# Longitudinal Live Imaging Derived 4D Hemodynamics and Dynamic Tissue Mechanics Across Outflow Tract Morphogenesis

**DOI:** 10.1101/2025.08.26.672464

**Authors:** Gening Dong, Jaehyun Rhee, Shivani Kumar, Molly E. Drumm, Henrik Lauridsen, Mahdi Esmaily-Moghadam, Jonathan T. Butcher

## Abstract

Growth and remodeling of the cardiac outflow tract (OFT) is poorly understood but associated with serious congenital heart defects (CHD). While only a minority of CHDs have identifiable genetic causes, the functional roles of mechanical forces in OFT remodeling are far less characterized. A key barrier has been the lack of longitudinal investigations examining the interplay between dynamic blood flow and wall motion across clinically relevant stages. Here, we developed a live high-frequency ultrasound derived 4D moving-domain computational fluid dynamics (CFD) simulation approach, enabling longitudinal quantification of OFT hemodynamics and tissue mechanics in the same embryos across Hamburger-Hamilton (HH) stage 21 to HH27. We found that rising wall shear stress (WSS) strongly correlates with tissue extension in the distal OFT, whereas the proximal OFT experiences increasing expansive strains and higher hydrostatic stress with heartbeats. Additionally, we identified a double-helical flow pattern in the OFT lumen, possibly reflecting an evolutionary legacy for directing oxidized and non-oxidized blood flow before great vessel septation. Together, our results advanced insights in how hemodynamic forces and tissue stress contribute to OFT remodeling and septation.

## INTRODUCTION

The cardiac outflow tract (OFT) is a rapidly growing and remodeling structure during embryogenesis, involved in several key morphological processes, such as aortic and pulmonary great vessels septation and semilunar valves formation (1–3). Abnormal remodeling of the OFT, such as the failure of septation or misalignment of the outlets with the ventricles, can lead to various congenital heart defects (CHDs) including persistent truncus arteriosus, double outlet right ventricle, and transposition of the great arteries (4, 5). CHDs are among the most diagnosed congenital disorders, which occur in 1-2% of live births worldwide and up to 35% of pregnancy loss. (4, 6–9). Malformations in the OFT are associated with about 30% of CHDs, causing significant morbidity and mortality clinically (4, 5, 8, 9). While genetic mechanisms of CHDs have been extensively studied, less than 30% of CHD cases have a genetic cause (6, 10). Abnormal mechanical stimuli such as hemodynamic perturbations and altered mechanical forces have been well recognized as alternative explanations of cardiac malformation (11–13). Recent studies demonstrate that hemodynamics forces are also essential for OFT cushion formation and valve remodeling in part through operating a variety of hemodynamic-responsive transcription factors and signaling pathways (14–17). Thus, a deeper understanding of hemodynamics and tissue mechanics during normal OFT morphogenesis will serve as a baseline for identifying cause and progression of CHDs.

The small size, rapid dynamic remodeling, and tortuous anatomy of the OFT pose significant challenges to quantitatively analyze biomechanical phenomena. Computational fluid dynamics (CFD) have thus been widely used to quantify blood flow within OFTs. Early computational studies used various imaging techniques, such as micro-CT and confocal microscopy, to acquire the *in vivo* geometry of embryonic OFT for image-based CFD simulations (18, 19). These studies used static models from fixed tissue, resulting in limited information on the spatiotemporal mapping of hemodynamics and tissue strains. More recent studies emphasized the interaction between cardiac wall motion and hemodynamics using 4D (3D over time) models generated from optical coherence tomography (OCT) in avian models (20–22). Due to low penetration depth of OCT, these studies only focus on early cardiac looping stages (HH13-18) prior to the formation of the OFT cushions (23). Other studies used inverted epifluorescence microscopes, confocal laser scanning microscopy, or light-sheet fluorescence microscopy (LSFM) to build 3D (2D over time) or 4D models of zebrafish embryos (24–27). However, zebrafish heart is a two-chambered organ that lacks the separation of oxygenated and deoxygenated blood in a four-chamber heart found in mammals and avians. As a result, zebrafish model is not suitable for the investigation of OFT septation and remodeling, whereas optical imaging techniques are not readily applicable to non-transparent animal embryos with four-chambered hearts. As a clinically extensively utilized non-invasive imaging tool, high-frequency ultrasound provides comparable temporal resolution, sufficient spatial resolution, and higher penetration depth to capture the cardiac wall motion at later stages, as demonstrated by studies in both OFT and chambers (28–30). While the blood flow dynamics and wall motions have been increasingly explored, a limited number of studies have performed analysis across development. Recent quantitative studies on growth and remodeling of embryonic cardiovascular structures across stages relied on fixed tissue imaging or 3D static models (18, 31– 33), whereas 4D dynamic modeling based on OCT provides information only on earlier cardiac looping stages before gestationally survivable but clinically serious CHDs arise (34). Moreover, these cohort studies examined developmental blood flow dynamics using pooled embryos at the population level, which is susceptible to bias due to batch effects and embryo-to-embryo variation. Similarly, far less is known about strain patterns in the OFT during morphogenesis. Experimental approaches have demonstrated that cyclic strain plays a role in mediating endothelial-mesenchymal transformation (EMT) in early heart valve morphogenesis (35), stimulating myocardial wall development and maturation (36), and is correlated with growth in embryonic cardiovascular development (37, 38). Therefore, quantitative analysis correlating fluid shear stresses, cyclic strains on cardiac wall, and anatomic geometry changes across development are needed to understand the interaction between mechanical forces and morphogenesis in the embryonic OFT. Together, these knowledge gaps highlight the absence of longitudinal investigations of dynamic blood flow and tissue mechanics across growth and remodeling in a four-chambered heart, which will significantly enhance experimental sensitivity for identifying developmental patterns.

Thus, the objective of this study is to quantify spatiotemporal mapping of the hemodynamic and dynamic tissue strains driving OFT remodeling in a four-chambered chick model due to its low cost, easy access, similar cardiac structure and developmental process to human embryo via *ex ovo* high-frequency ultrasound-derived longitudinal 4D moving-domain CFD simulations across CHD emerging developmental stages.

## MATERIALS AND METHODS

### Chick embryo culture and preparation

Fertilized white Leghorn eggs were incubated blunt end up in a continuous rocking incubator at 37.5 °C and approximately 50% humidity for 72 hours. On the 3rd day, the embryos were moved to an *ex ovo* culture system as previously described (39). The ex-oxo-cultured embryos were placed back into the incubator until they reached the desired stages (HH21 - HH27). Before imaging, embryos were transferred into a portable incubator at 37 ± 0.5 °C along with two 9 oz cups filled with distilled water to keep the appropriate humidity.

### Ultrasound imaging and image process

Imaging was performed using the Vevo 2100 high-frequency ultrasound system (Visual Sonic Inc., Toronto, Canada) with a 30-70 MHz MS700 transducer. During scanning, a 37 ± 0.5 °C water bath (IVYX Scientific Inc., Seattle, USA) and a heat lamp were used to control the temperature of the embryos. At least 1 ml of warm Tyrode-HEPES solution was placed between the embryo and the ultrasound transducer head for coupling. During scans, embryos were consecutively moved along the scan plane’s z-axis with a step size of 30 - 50 μm until the entire cardiac OFT was imaged. At each z position, a spatiotemporal (2D+T) image containing at least 15 cardiac cycles was captured at 130 frames per second in B-mode guided ultrasound. After imaging, the embryos were placed back into the incubator and allowed to recover until they reached the next desired stages.

The image processing technique was developed using MATLAB (The MathWorks Inc., Natick, USA) based on a quadratic ensemble averaging algorithm to average the time stack into one full cardiac cycle to minimize temporal error and increase the contrast between tissue and blood spaces. 3D reconstruction was performed based on previously established protocol (40). Shortly, images with the same phase of the cardiac cycle were first identified by calculating the correlation value among images:

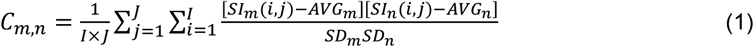

where *C*_*m,n*_ was the correlation value between the image at the *m*^*th*^ and *n*^*th*^ time frame, *SI*_*k*_(*i, j*) was the signal intensity of the pixel at coordinates (*i, j*) in the image at the *k*^*th*^ time frame, *AVG*_*k*_ and *SD*_*k*_ were the mean and standard deviation of pixel intensity of the image at the *k*^*th*^ time frame, respectively, and *I* and *J* are the total numbers of pixels in x and y directions.

The peak correlation values were taken as matching cardiac phases and used to compute the quadratic average *Q*(*AVG*) for all correlated frames using the following equation:

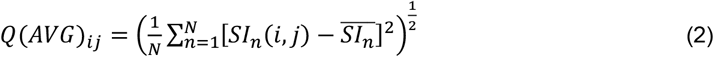

where *N* was the total number of matching frames at one cardiac phase, *SI*_*n*_(*i, j*) was the signal intensity at coordinates (*i, j*) of the image within the *N* matching images, and 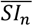 was the arithmetic mean of *SI*(*i, j*) over all images to be averaged.

For synchronization of different slices, mid-ventricular slice with easily recognizable anatomical structures was first selected as the reference slice. Cross-correlation values between two adjacent slices after ensemble averaging were calculated using formula (1). The ensemble averaged one cardiac cycle from the neighboring slices were adjusted to match the cardiac phases with the reference slice based on identified peak correlation values.

### Dynamic reconstruction

An automatic boundary-tracking technique was utilized to track the motion of the cardiac wall for dynamics reconstruction.

The endocardial layer of the cardiac OFT was first segmented at a time frame with easily recognizable anatomical structures using the automated segment method based on pixel intensity in the open-source software, 3D Slicer (41, 42). The frame where segmentation was performed was then used as the fixed and reference frame for the following sequence registration. Non-rigid registration based on third-order B-Spline interpolation with custom parameters was performed to the volume sequence at each stage using the plugin module, Sequence Registration (41), to track the endocardial boundaries. The resulting transformation sequence was then applied to the segmentations to obtain dynamic 4D geometries.

### Moving-domain simulation

4D moving domain CFD simulations were performed using Multi-Physics Finite Element Solver (MUPFES) (43–45). For mesh generation, segmentations were smoothed using an open-source 3D mesh processing software, MeshLab (46). Laplacian smoothing was applied globally to remove artifacts due to discrete acquisition in z steps. Smoothed models were then meshed into 400,000 - 850,000 tetrahedral elements with 50,000 surface elements on the surface using Tetgen (47).

The previously described boundary tracking algorithm obtained wall boundary motion. A Fourier transform was performed on the displacement functions of surface mesh nodes to remove the artificial oscillations due to discrete acquisition in time, where high-frequency noise was filtered out. Smoothed wall motion was then input as wall motion boundaries (Dirichlet boundary condition) to the CFD solver. An isotropic linear elastic model with a Poisson’s ratio of 0.49 (48) was used for the cardiac wall, where no-slip boundary conditions were applied.

Simulations were conducted with a time step size of 0.05 ms, and a complete cardiac cycle was interpolated into 8,000 - 17,000 time steps, depending on the heart rate. The blood was treated as incompressible and Newtonian fluid with a constant density of 1,040 kg/m^3^, and a viscosity of 3.5 × 10^−3^ Pa · s. Neumann boundary conditions were applied to the inlet and outlet surfaces. Time-varying pressure profiles were used as the mid-ventricular inlet boundary conditions. Ventricular pressure traces at HH21, HH24, and HH27 were adopted from previously published papers (49– 51). Vascular resistance was used as the distal outlet boundary conditions for the HH21 and HH24 simulations, whereas the outlet boundary conditions were set free for the HH27 simulation. Previously published resistance values were assumed constant throughout a cardiac cycle (52).

During the cardiac phases when the distal cushions or the proximal cushions are shut, the wall motion was not modeled due to the limitation that intersective mesh elements can lead to wall leaking and non-convergence in the simulation. To simulate flow blockage during cushion shut, elements in the cushion regions were selected and extracted as separated boundary surfaces, to which zero flow rate boundary conditions were applied during the corresponding time points when the cushions shut. Time frames when the cushions shut were determined by the live ultrasound cine-images of each embryo at each stage.

### Simulation results post-processing

An open-source visualization application, Paraview (53), was used for simulation results post-processing. Simulation results were saved every 50 time steps (2.5 ms). For each element, the oscillatory shear index (OSI) was calculated by:

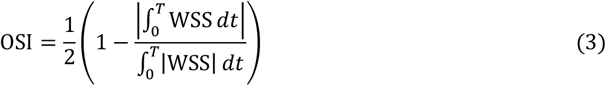

In addition, normal helicity (*h*) was computed to learn more about the flow pattern in the OFT during morphogenesis, which is defined as:

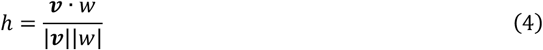

where *v* is the velocity vector, and *w* is the vorticity.

For surface elements, Green-Lagrangian strain tensor was computed based on the nodal displacements as the local strain, which is defined as:

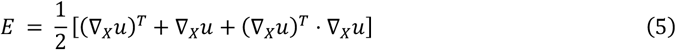

where **E** was the Green-Lagrangian strain tensor, *x* was the coordinate vector, and **u** was the displacement vector from the time frame where the model reaches its minimum volume. Principal strains σ_1_, σ_2_, and σ_3_ were then used for analysis by calculating the eigenvalues of the strain tensor, where we set:

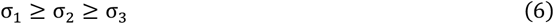

Laplace’s law was used to estimate circumferential stress using the following formula:

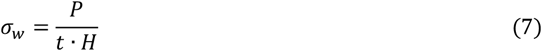

where P was the blood pressure, *t* was the wall thickness, and *H* was the mean curvature, which is defined as:

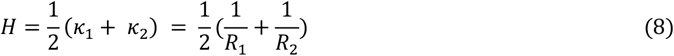

where *k*_1_ and *k*_2_ are the principal curvatures, and *R*_1_ and *R*_2_ are radii corresponding to the principal curvatures.

### Statistical analysis

Hemodynamic indices, tissue strains, and morphological changes were compared quantitatively among regions and developmental stages when possible. Results were summarized in means and standard deviations over a specific region or over a cardiac cycle. Statistical comparisons were made using ggplot2 package v3.4.4 (54). Paired t-tests were used when comparing means among developmental stages, while unpaired t-tests were used when comparing among regions. P<0.05 denotes significance.

## RESULTS

### Modeling of the cardiac outflow tract

4D high frequency (50 MHz) ultrasound imaging was performed on a total of 4 embryos on three consecutive days from the 4th day (D4) to the 6th day (D6) after incubation, during which the embryos were at Hamburger-Hamilton (HH) stage 21, HH24, and HH27, respectively. Cine-images of the embryos were acquired over 30 - 55 transverse planes spaced 30 - 40 μm with a temporal resolution of 130 frames per second (Fig. 1A). The heart rate (HR) of each embryo was monitored during imaging to confirm embryo viability during the multi-day ultrasound scan. The embryo’s HRs are consistent with earlier population studies (52, 55), demonstrating the embryo’s viability was not affected by the multi-day ultrasound scans (Fig. 1B). Each cine-image was reconstructed into one full cardiac cycle with 55 – 70 distinct cardiac phases with increased signal-to-noise ratio based on a quadratic ensemble averaging algorithm (Fig. 1C). Post-processing images have a spatial resolution of 7 – 9 μm/pixel in the scan plane and 30 – 40 μm voxel depth. To correct the oscillatory artifacts due to discrete acquisition in space and in time, Laplacian smoothing and Fourier transform was performed to the meshed segmentations and the displacement functions, respectively (See Materials and Methods). Model volumes were examined before and after smoothing, showing a satisfactory match within cardiac cycle and no difference statistically (Fig. 1 D, E).

**Figure 1.**
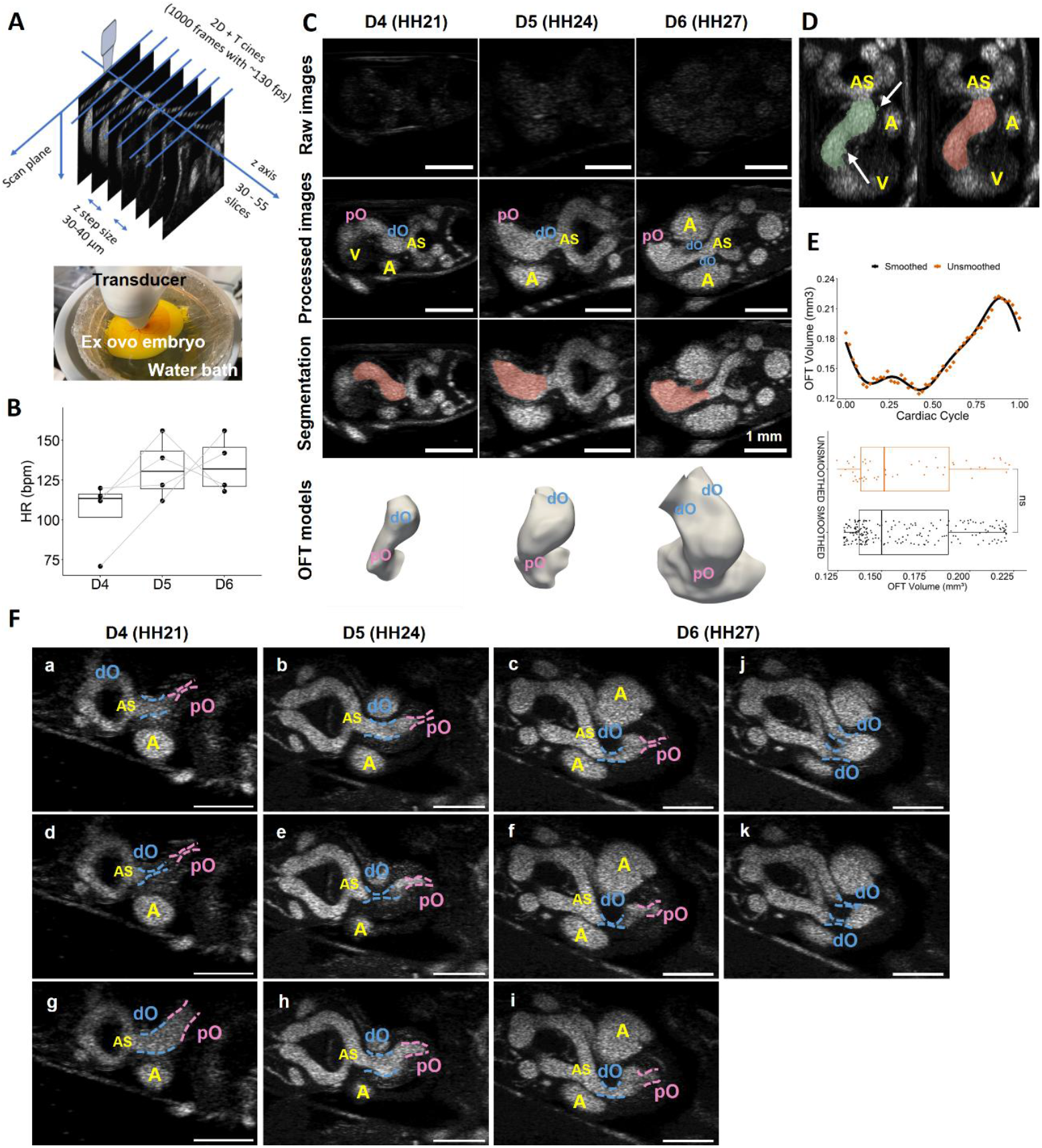
Longitudinal live ultrasound imaging-derived dynamic model reconstruction. **A)** Schematic of 4D ultrasound scan. **B)** Heart rate of scanned embryos, demonstrating the embryo’s viability was not affected by the multi-day ultrasound scans. Data points from the same embryo are linked with gray line. **C)** Raw ultrasound cine images (first row), processed images (second row), smoothed segmentation overlapping on processed images (third row), and the models of the OFT (last row, containing part of the ventricles as inlets) at HH21, HH24, and HH27, respectively. **D)** Unsmoothed segmentation (left), and smoothed segmentation (right) of a D4 ultrasound dataset. **E)** Volume of a D4 OFT obtained from the unsmoothed segmentation of ultrasound stacks and from smoothed reconstructed dynamic models over a cardiac cycle, showing no difference. **F)** Transition in cushion motions across development. At all stages studies, pO cushions close prior to the dO cushions (a-c). At HH21, the entire OFT lumen was able to fully collapse (d), and the dO and pO opened at the same time (g). At HH24 and HH27, the pO cushions open prior to the dO cushions (e, f, h, i). In addition, at HH27 when the dO separates, two pairs of dO cushions close and open simultaneously (j, k). OFT, outflow tract; dO, distal outflow tract; pO, proximal outflow tract; A, atria; V, ventricles; AS, aortic sac.

### Morphological changes from 4D modeling

To provide a better understanding of the hemodynamic and anatomic changes during the critical stages when heart valves start to form, we reconstructed the cardiac motion in full-scale chick OFT geometric models based on the cine-images of the same embryos at 4 (HH21), 5 (HH24), and 6 (HH27) days old.

Reconstructed 4D ultrasound image stacks -support that OFT cushions in early embryonic hearts served as dynamic valves as the pO and dO regions undergo periodical luminal collapse. At HH21, the pO and dO were connected and the OFT lumen was able to close completely, where at later developmental stages, vertically separating dO and pO were observed, and a bulging plenum region appeared in between (Fig. 1 C), which most likely corresponds to the intermediate OFT or the dog leg bend as previously described (56, 57). Though expansion and contraction were also observed in the intermediate OFT, no full luminal collapse occurred. The formation of the intermediate OFT was accompanied by a dramatic transition in the physical cushion action. At HH21, the pO and dO regions recover from the lumen collapse simultaneously despite the pO cushions closing first (Fig. 1F, Video S1A). At later stages after the intermediate OFT appearance, the pO and dO cushions function at different phases of cardiac cycle during both closing and opening, following a cushion motion pattern of pO closing, dO closing, pO opening, and dO opening (Fig. 1F, Video S1B). Notably, at HH27 when the dO has separated into two outlets, two pairs of dO cushions open and close simultaneously (Fig. 1F, Video S1C). Due to mesh singularity constraints, the total lumen collapses in the pO and dO when the cushions are fully closed were not modeled using wall motion. Instead, to simulate the luminal collapse at the proximal and distal cushions, zero inlet flow rate was applied to the mesh nodes located in the cushion regions during the time steps when the cushions were supposed to be closed according to the live ultrasound cine-images. Further, we synchronize the cardiac phases among embryos by aligning the time point when the pO fully shut.

We then examined the volume change of the OFT both within the cardiac cycle at the same stage, and at a specific cycle phase across development. The OFT volume was approximated by summing up the volumes of tetrahedral elements in the reconstructed mesh. Only elements located in the OFT region were selected to exclude the mid-ventricular inlets and distal vascular outlets. Volume normalized to the minimum volume at the time point when the cushions shut was measured within a cardiac cycle (Fig. 2 A). Though no difference was found in the mean volume of the OFT, most embryos showed volume decreases from HH21 to HH24 and from HH24 to HH27 (Fig. 2 C). OFT stroke volume, defined as the difference between the maximum volume and the minimum volume within a cardiac cycle, increased significantly from HH21 to HH27 (Fig. 2 B). This was accompanied by a significant increase in the OFT expansibility (Fig. 2C), leading to increased normalized volume change within heartbeats (Fig. 2 A). When looking at individual embryos longitudinally, the stroke volume and OFT expansibility showed the same developmental patterns. Half of the embryos’ stroke volume decreased from HH24 to HH27, which is consistent with the HR change that 2 out of 4 embryos’ HR decreased from HH24 to HH27 (Fig. 1B). This is likely due to a slight difference in temperature during ultrasound scan, suggesting a correlation between embryo HR and OFT stroke volume. Further, we examined the surface area change within a heartbeat during OFT development. Like volume change, statistically significant increases in OFT surface area stretchability were observed from HH21 to HH27 without any significant changes in the time-average surface area across development (Fig. 2 D).

**Figure 2.**
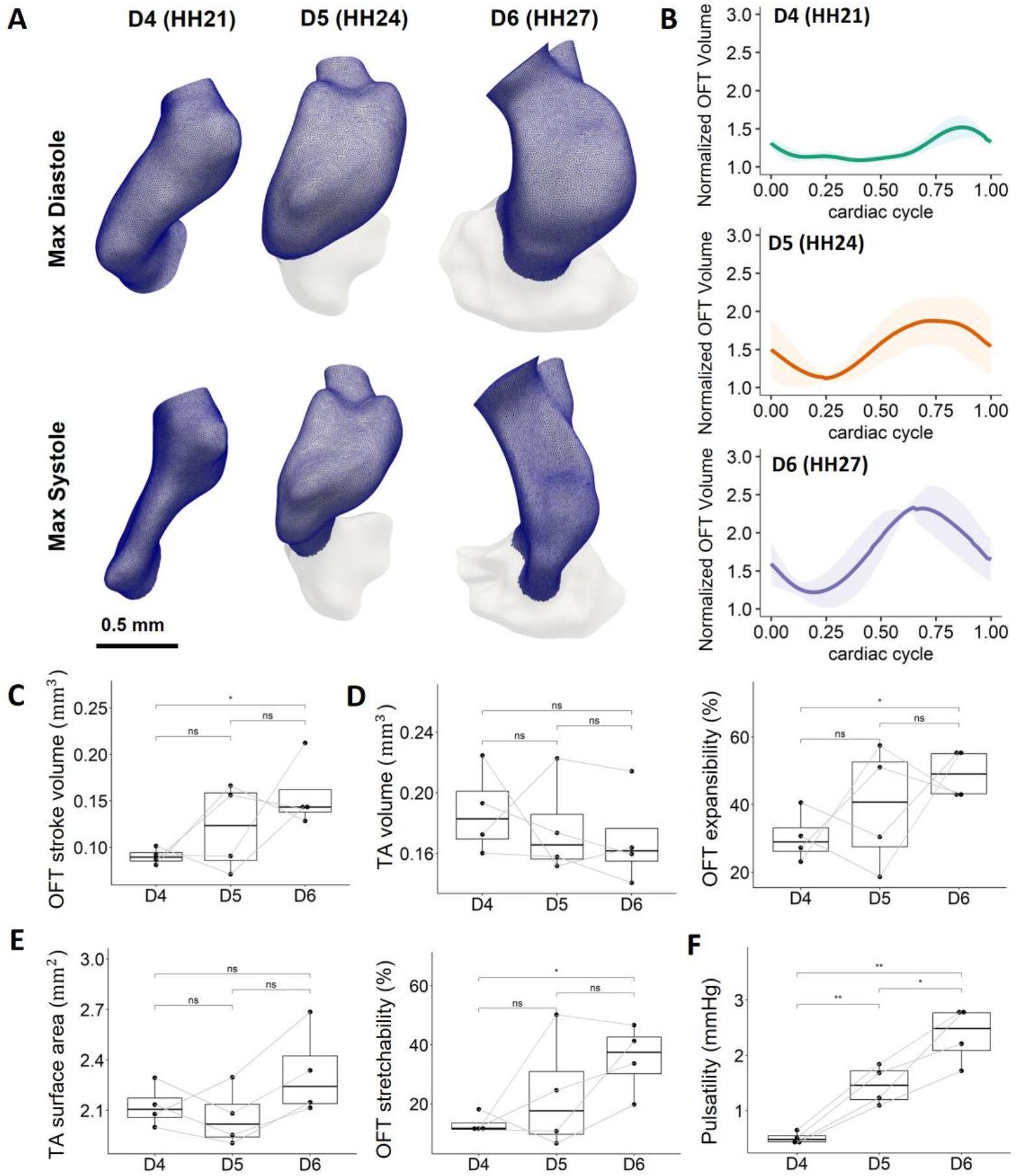
Anatomic and cardiac function changes during cardiac OFT morphology. **A)** The reconstructed OFT models were meshed into tetrahedral elements. Volumes of elements located in the OFT region were summed up to approximate the OFT volume within a cardiac cycle. **B)** Normalized volumes of the OFT within a cardiac cycle at different developmental stages. Volumes were normalized to the minimum volume when the cushions shut. Data is shown in mean ± SEM (N = 4). **C)** OFT stroke volume was found to increase significantly from D4 to D6. **D)** No difference in TA volume of the OFT over development, however, the volumetric expansibility increased significantly from D4 to D6. **E)** No difference in TA surface area of the OFT over development, but the surface area stretchability increased significantly from D4 to D6. **F)** The pulsatility of the OFT increased significantly from D4 to D6. SEM, standard error of the mean; TA, time-averaged. OFT stroke volume = max V – min V (within a cardiac cycle); Pulsatility = max pressure – min pressure (within a cardiac cycle); OFT expansibility (%) = [max V – TA V]/TA V * 100%; OFT stretchability (%) = [max area – TA area]/TA area * 100%. Data points from the same embryo are linked with gray line. *p <= 0.05, **p <= 0.01, ***p <= 0.001,****p <= 0.0001.

### Local *in vivo* Hemodynamics

We developed a live image-based 4D moving-domain simulation strategy, which enabled the quantitative study of the hemodynamic indices within a beating embryonic OFT across developmental stages. Time-varying wall motion based on boundary tracking was applied to the corresponding OFT segmentation. Coordinates of surface mesh nodes were extracted at every time frame and reorganized into displacement information of every mesh node after removing high-frequency noise (see Materials and Methods). Nodal displacements at every time frame were then input as an additional boundary condition to the mesh wall. By considering the interaction between time-varying wall motion and hemodynamics, we provided new insights into spatiotemporal variations of hemodynamic indices and tissue strains.

Pulsatility of the OFT lumen, which was defined as the difference between the maximum and minimum pressure within a cardiac cycle, was first examined. Pulsatility increased significantly across development, and all embryos showed constant pulsatility increases from HH21 to HH27 (Fig. 2 E). Taking together with the increasing expansibility and the stretchability over development, these could suggest that OFT tissue may withstand higher pressure difference via greater cardiac motion as damping.

Moving-domain CFD simulation results also showed significant temporal and spatial variations in WSS magnitudes (Fig. 3A, S4, Video S2A-C). Peak WSS values were observed in regions with smaller diameters and higher velocities such as the inner and outer curvature, the distal outlets, and the root of distal vascular bifurcation at HH27, generally consistent with locations where pO and dO cushions are located (Fig. 3A, S1A). Spatially averaged WSS magnitudes in each region showed a significant increase in the dO and in the intermediate OFT with development, whereas no difference was found in the pO from HH21 to HH24 (Fig. 3B). Looking at individual embryos longitudinally, all 4 embryos studied showed constant increases across development in both peak and time-averaged WSS in the dO, whereas WSS in the pO decreased from HH21 to HH24 in 2 of 4 embryos (Fig. S3A, B). When comparing between different regions, pO experienced higher WSS than dO at HH21 and HH24. However, dO started to experience significantly higher WSS than pO at HH27. This is consistent with significant increases in peak WSS in the dO from HH24 to HH27 (Fig. S3A). The intermediate OFT experienced the lowest WSS during stages after its formation at HH24 (Fig. 3B), which is likely a result of its lower velocities and vortex flow during cushion shut (Fig. S1A, B). This is consistent with the oscillatory shear index (OSI) results, that the peak OSI magnitudes are located at the intermediate OFT at HH24 and HH27 (Fig. S2A). In addition, higher OSI magnitudes also appeared at the cushion regions in the dO and pO, which is likely due to the cyclic closure and opening of the cushions. In addition, spikes of WSS can be seen at the opening of lumen after cushion shut (Fig. 3C), which was due to blood ejection from the ventricles through the OFT. An exception was observed in embryo 2 (E2) at HH27, where the OFT experienced peak WSS magnitudes at later cardiac phases. This could be a natural variation or a slight difference in developmental stage, as its pO experienced higher WSS than dO, which is a feature at younger stages in other embryos. Furthermore, the length of cushion closure time also varies among embryos, with the largest difference occurring at HH24 (Fig. 3C). However, this time difference decreased at HH27, indicating a possible transient from variance to robustness from HH24 and HH27.

**Figure 3.**
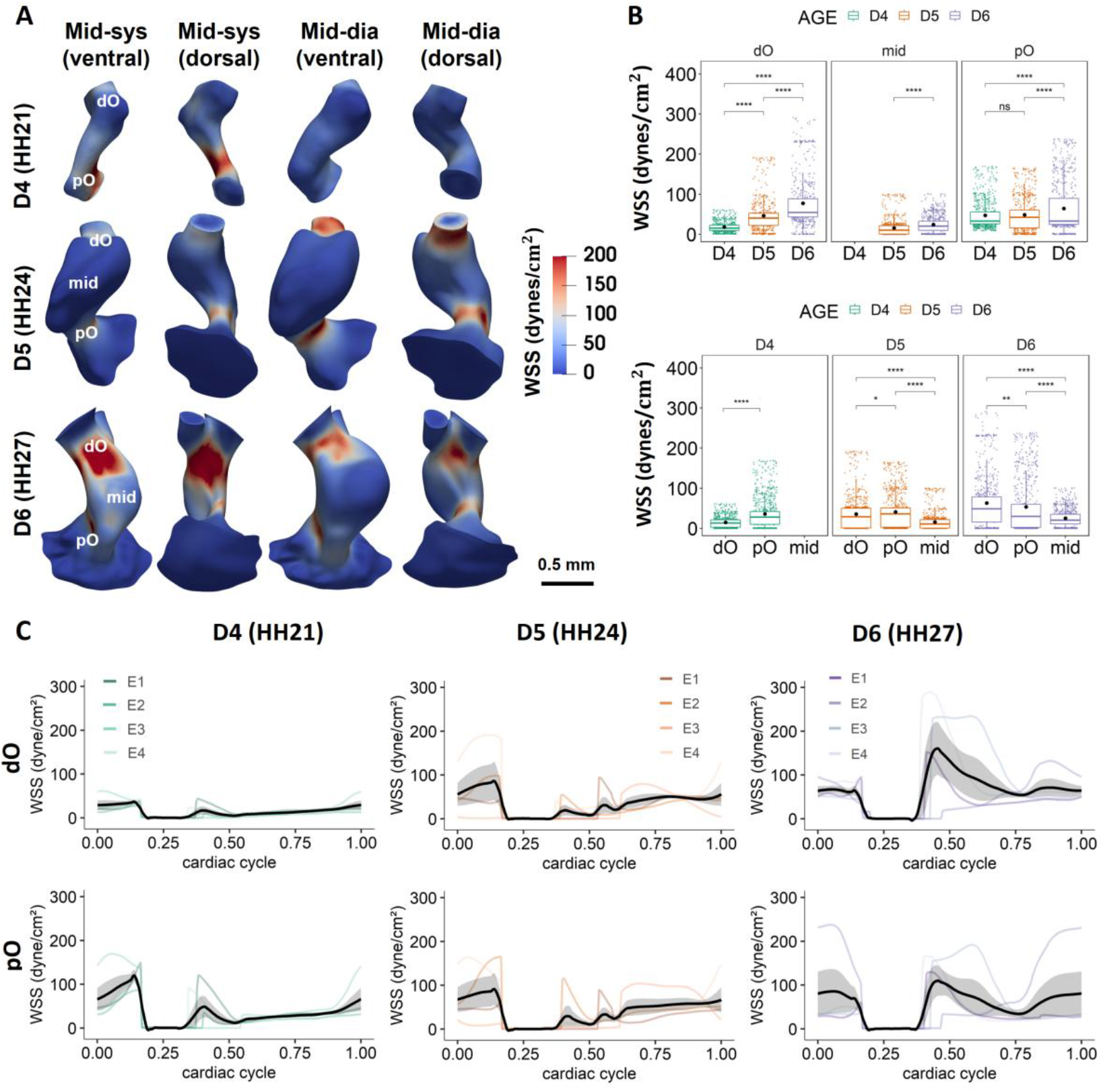
Regional WSS results. **A)** Time- and space-varying WSS magnitudes at different phases of cardiac cycle over development. Figures show the 3D renderings of the surface WSS magnitudes on the dynamic models of Embryo 1 at different developmental stages at mid-systole and mid-diastole. Higher WSS magnitudes locate at regions with smaller diameters such as the proximal cushions and distal cushions. **B)** Regional WSS magnitudes within a cardiac cycle increased significantly over development in the dO and in the intermediate OFT (mid). When comparing between different regions, the pO experience significantly higher WSS than the dO at HH21 and HH24. However, dO starts to experience significantly higher WSS at HH27. **C)** Mean WSS magnitudes in different regions over a full cardiac cycle across developmental stages for 4 embryos. Cardiac cycles were matched among different embryos based on the cardiac phase when the pO cushions shut. *p <= 0.05, **p <= 0.01, ***p <= 0.001,****p <= 0.0001.

### Local *in vivo* strain and stress

Further, we quantified the local tissue strains and stresses during OFT morphogenesis based on our live imaging-derived moving-domain CFD simulations. As the total lumen collapse was not able to be reconstructed using the finite meshes, the OFT model with a minimum volume within a cardiac cycle was used as the reference model to compute displacements of the cardiac wall. Thus, expansive strains were calculated based on nodal displacements from the cardiac phase when the OFT model reaches its minimum volume (see Materials and Methods).

Like WSS results, spatial and temporal variations were observed in strain magnitudes (Fig. 4A, S5, Video S3A-C). Development trends in spatially averaged strain magnitudes were found in the pO and the intermediate OFT, which experienced significantly larger strains across development (Fig. 4B). Though no statistical significance was found in the strain magnitudes at the dO region between HH21 and HH24, the dO experienced larger strains at HH27. These findings indicate larger cardiac wall movements at HH27, consistent with significantly greater expansibility and stretchability at HH27. Interestingly, cardiac tissue in the pO region experienced significantly higher strains than those in the dO at all stages examined (Fig. 4B), suggesting larger tissue deformations and wall motions in the pO during OFT remodeling. In addition, no spikes in strain were observed within a cardiac cycle, shown by generally arch-shaped smooth curves (Fig. 4C). This finding suggests that the OFT, like the ventricles, undergoes continuous cyclic contraction and expansion movements with heartbeats.

**Figure 4.**
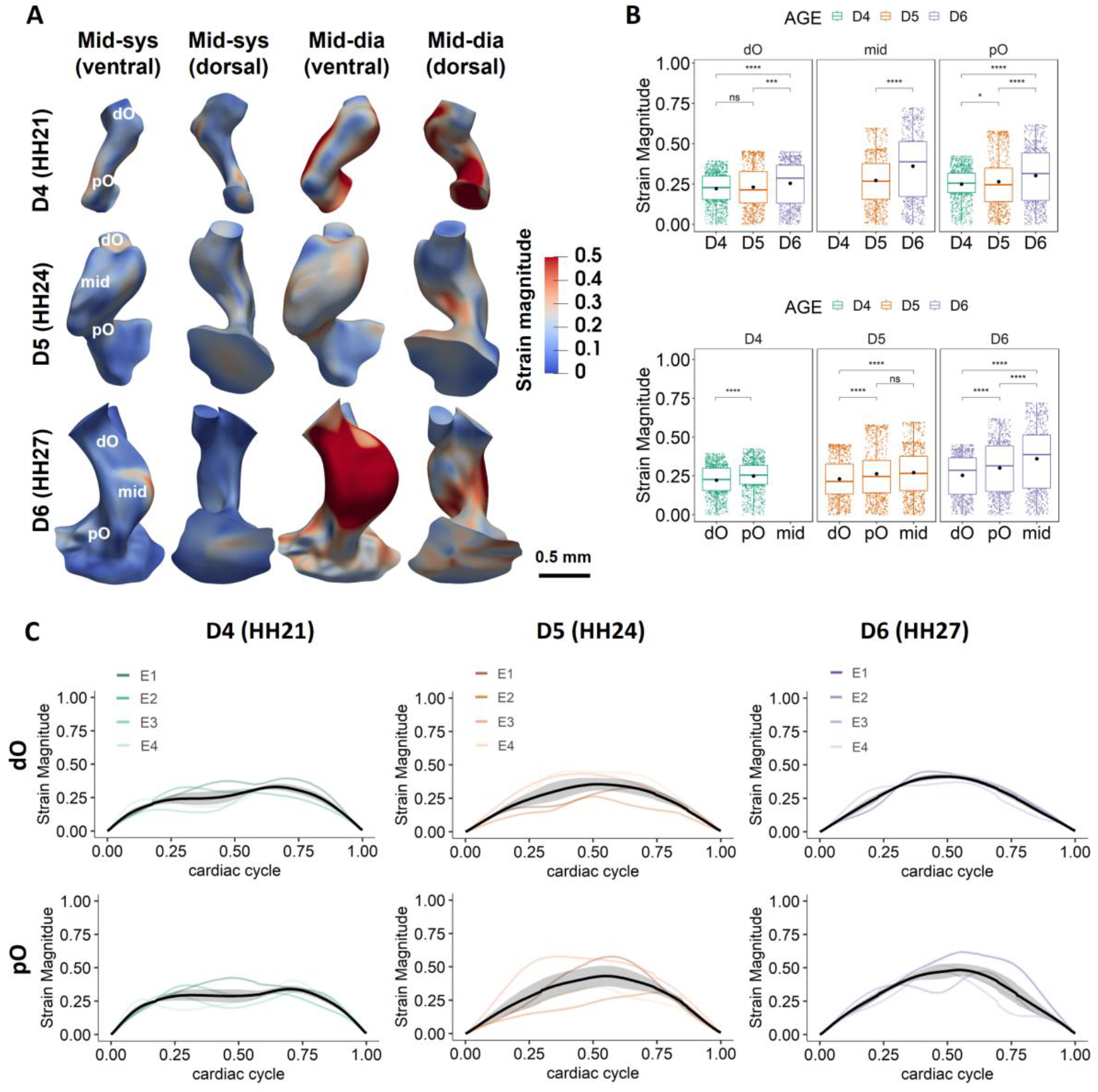
Regional strain results. **A)** The cyclic local strain magnitudes at different phases of cardiac cycle over 3 days. Figures show the 3D renderings of the cyclic strain on the dynamic models of Embryo 1 at different developmental stages at mid-systole and mid-diastole. Strains were calculated based on nodal displacements from the cardiac phase when the OFT model reaches its minimum volume. **B)** Plot shows mean strain magnitudes in different regions within a cardiac cycle over development. The pO experienced significantly higher strains with development. When comparing between different regions, the pO experienced significantly higher strains than the dO at all stages. **C)** Mean strain magnitudes in different regions over a full cardiac cycle across developmental stages for 4 embryos. Cardiac cycles were matched among different embryos based on the cardiac phase when OFT model reaches its minimum volume. *p <= 0.05, **p <= 0.01, ***p <= 0.001,****p <= 0.0001.

To understand the tensile and compressive stress patterns in the developing OFT, hydrostatic stress was calculated using Young-Laplace equation (see Materials and Methods). In regions where cushions are located, tissue experienced generally compressive stress (Fig. 5A, S6). An exception was observed in the outer curvature at HH21, where tissue experienced tensile stress during diastole. The inner curvature, as expected, was consistently under compressive stress due to limited change of curvature across cardiac cycle. Tensile stress was generally observed in the intermediate OFT. However, transfer between compressive and tensile stress was also observed in this region within a cardiac cycle, which is likely a result of its large cardiac motion and dramatical geometry changes with heartbeats. Statistical analysis showed that tissue in the pO experienced significantly higher stress magnitudes than that in the other regions, whereas tissue in the dO was under lowest stress when comparing among regions (Fig. 5B). These findings were in line with the pressure drop through the OFT lumen, that the pO experiences higher pressure than the dO. Spikes in stress magnitude were observed mostly during the cushion closure, when the pressure in the OFT boosted. Interestingly, significantly higher stress magnitudes were found in the OFT at HH24 than HH27 (Fig. 5B, C), despite the ventricular ejection pressure at HH27 is higher than that at HH24 with embryonic development, which might be a result of larger lumen diameter at HH24.

**Figure 5.**
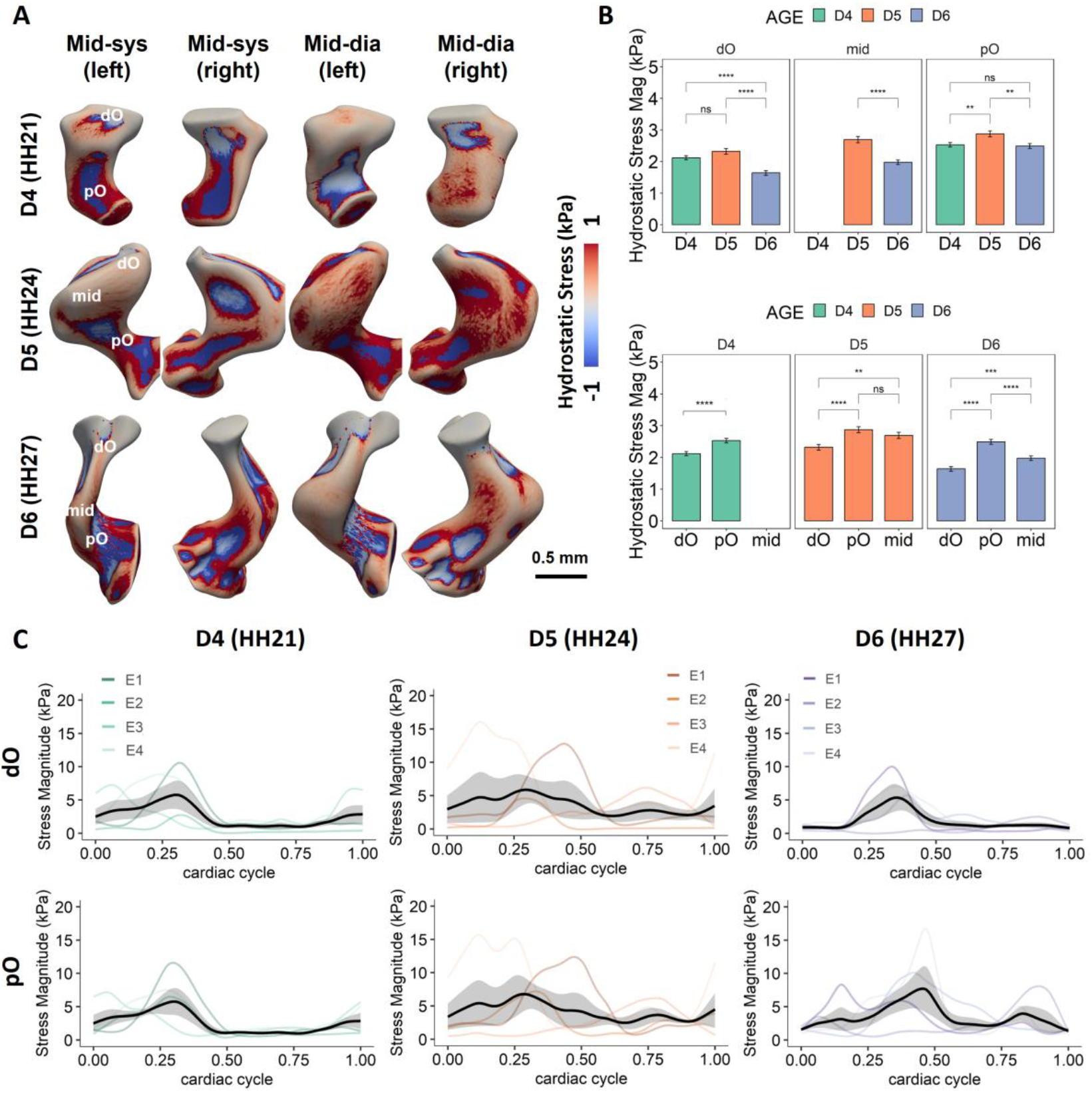
Regional hydrostatic stress results. **A)** The cyclic local hydrostatic stress at different phases of cardiac cycle over 3 days. Figures show the 3D renderings of the wall stress on the dynamic models of Embryo 1 at different developmental stages at mid-systole and mid-diastole. Positive values indicate tensile stress, whereas negative values indicate compressive stress. **B)** Plot shows mean stress magnitudes in different regions within a cardiac cycle over development. Embryos at HH24 were found to experience significantly higher wall stress than those at HH21 and HH27. When comparing between different regions, the pO experience significantly higher stress than the dO at all stages. **C)** Mean wall stress magnitudes in different regions over a full cardiac cycle across developmental stages for 4 embryos. Cardiac cycles were matched among different embryos based on the cardiac phase when the pO cushions shut. *p <= 0.05, **p <= 0.01, ***p <= 0.001,****p <= 0.0001.

### Hemodynamic force and strain driven OFT development and remodeling

To understand if and how hemodynamic force and tissue mechanics contribute to the remodeling of the developing OFT, we first examined the spatial correlation between local WSS and tissue strain, where elemental time-averaged strain and WSS magnitudes were used for correlation determination. Correlation analysis yielded statistically significant (p<0.05) but negligible correlations (R<0.1) at all developmental stages examined (Fig. S3F-H). Consistently low correlation coefficients suggest no meaningful correlation between WSS and tissue strain, indicating that WSS and tissue mechanics are not morphologically related. We further correlated the WSS and tissue strains with the OFT volume expansibility and surface area stretchability. Developmental change in time-average wall shear stress (TAWSS) showed significant positive correlations with OFT stretchability in both dO and pO, whereas no statistically significant correlation between WSS and OFT expansibility was determined (Fig. 6A). This analysis revealed that hemodynamic forces play a significant role in driving tissue stretchability growth in both dO and pO. Interestingly, stretchability was positively correlated with tissue strains in the dO as well (Fig. 6B). This suggests that strain induced by wall motion may be a secondary driver of stretchability development in dO following local WSS. In addition, dO showed a positive tissue strain-OFT expansibility trend with a strong correlation (Fig. 6B). This is also demonstrated by similar developmental trends in OFT expansibility to those of dO peak and time-averaged strain (Fig. S3D, E). These results suggest cyclic strain is strongly correlated with both tissue expansibility and stretchability development in distal OFT remodeling. Though no significant trend was identified for strains and volumetric expansibility or surface area stretchability in the pO, increasing regional tissue strains associated with increasing volume more than with surface area changes, demonstrated by higher correlation coefficient (R) in the pO region (Fig. 6B).

**Figure 6.**
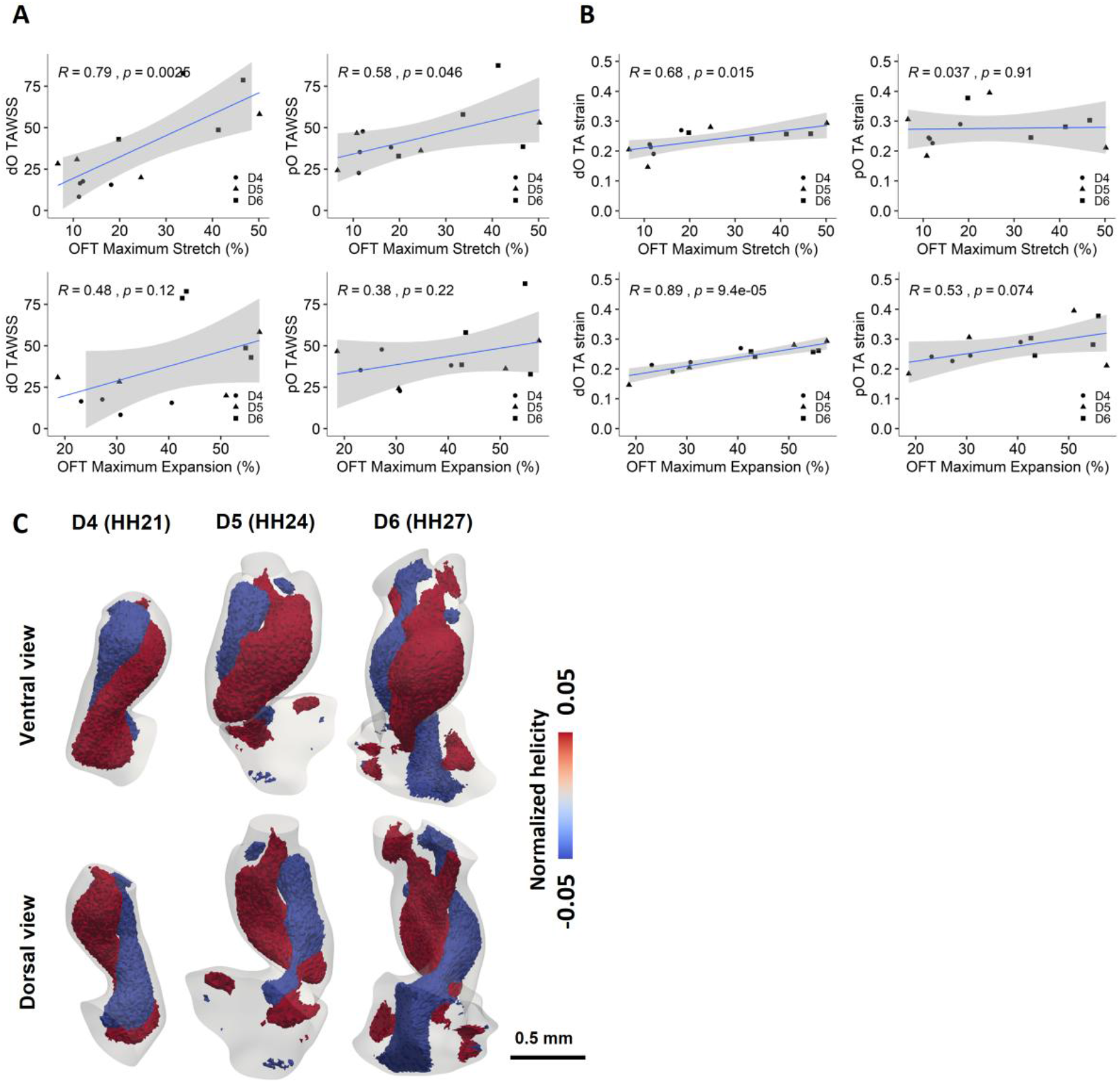
Hemodynamic force and strain driven OFT development and remodeling. **A)** Linear regressions were plotted to identify correlations between hemodynamics and surface area or volume changes. TAWSS and OFT stretchability are strongly correlated in both the dO and pO, but only weak correlations between TAWSS and OFT expansibility in all regions. **B)** Strong correlation between TA strain values and OFT expansibility in dO, TA strain and OFT stretchability correlate more in pO, shown by higher correlation coefficient (R). **C)** Double-helical flow pattern observed during blood ejection after dO cushions opening, shown in the models of Embryo 1 at different developmental stages. Red, elements with normalized helicity > 0.05 (clockwise rotation); blue, elements with normalized helicity < -0.05 (counterclockwise rotation). OFT Maximum Stretch (%) = [max SA – TA SA]/TA SA * 100%; OFT Maximum Expansion (%) = [max V – TA V]/TA V * 100%. TA, time averaged; V, volume; SA, surface area.

To further uncover the hemodynamic driven OFT remodeling, normalized helicity was calculated in the developing OFT during forward ejecting blood after dO cushion opening. In forward blood flow from the ventricles to the aortic arches, positive helicity indicates a clockwise rotating flow, whereas negative helicity indicates a counterclockwise rotating flow (58). A double-helical flow was observed at all stages studied over development, with two counter-rotating flows twisting around each other (Fig. 6C, S7). This flow pattern formed during blood ejection when dO cushions open and was observed from the ventricular inlet towards the distal vessels. The twisting was more significant at the earlier development stage than at later stages. In addition, the double-helical flow is more significant in the proximal region at HH27, where interactions of the two counter-rotating flows increased in the distal region.

## DISCUSSION

Significant efforts have been made to quantitatively characterize hemodynamics and tissue mechanics during normal cardiac morphogenesis. Previous studies have provide valuable insights using imaging-derived simulations based on static geometries (11, 18, 59), 2D moving-domain models (25), mathematically modeled wall motion assumptions (29), 4D models at certain early stages (20, 21, 28, 29), or population-level analysis (1, 32, 34, 59). However, a gap in direct and longitudinal approaches to resolve the interplay between blood flow and tissue mechanics in individual embryos across developmental stages has hindered efforts to uncover the mechanobiological phenomena that shape the morphing heart. We here performed the first longitudinal 4D flow simulations within a beating embryonic OFT in the same embryos based on *in vivo* imaging derived cardiac motion, which enabled the quantitative monitoring of the hemodynamic indices and dynamic tissue stress across clinically relevant developmental stages.

We advanced swept-source high-frequency ultrasound scanning techniques without the usage of polyurethane membrane or toxic ultrasound gel to increase embryo survival rate. We also integrated *ex ovo* chick culture with ultrasound scan for larger tissue access. Together, these techniques enable direct and longitudinal tracking of the same embryos across stages. Increase in HR from D4 to D6 supports embryo growth despite multi-day ultrasound scans. Although 2 embryos showed a slight decrease in HR from D5 to D6, this could be natural variations or a slight difference in temperature or humidity during scan. Stable HRs during the three consecutive days of imaging demonstrate our imaging strategy can capture dynamic pictures of cardiac morphogenesis in the same chick embryos across development without embryo viability being affected. Notably, our longitudinal studies were performed at clinically relevant stages, where CHDs that require surgical interventions and lead to high fetal or neonatal death arise. Tetralogy of Fallot (ToF) and transposition of the great arteries (TGA) are reported to be the most and second most common critical CHDs in live births, respectively (60). In addition, pregnancy loss occurred at 24-40 weeks gestational age due to diagnosed CHDs have been reported to be larger than 5%, among which the highest incidence of fetal demise was related to valve stenosis, double-outlet right ventricle, and truncus arteriosus (9, 61). These clinically relevant CHDs due to defective septation or structural maturation are highly associated with abnormal OFT remodeling during the development stages we studied. On the other hand, earlier defects in cardiogenesis, such as genetic defects, heart tube formation defects, and defective cushion remodeling during cardiac looping, result in early embryonic lethality that limited clinical intervention can be done (62–65). This emphasizes the importance of understanding hemodynamics and tissue strains across clinically relevant later stages of OFT growth and septation.

Longitudinal ultrasound scanning technique enables *in vivo* monitoring of anatomy and tissue wall motion across development. The reconstructed 4D geometric models showed that the time averaged OFT volume did not change significantly during morphogenesis despite the presence of an expanding intermediate OFT region at HH24 and HH27 that doesn’t fully collapse during cardiac cycle. Instead, most embryos showed a decreasing trend in time-average volume across development. This may be related to the high levels of cell apoptosis found in both the pO and the dO in earlier studies (66). Interestingly, the stoke volume increased significantly from HH21 to HH27 despite the reduce in volume, suggesting the OFT is able to transport blood flow from the ventricular ejection more effectively with a smaller volume. In addition, both maximum volumetric expansibility and surface area stretchability increased significantly across development, suggesting the tissue properties change jointly with tissue motions. The increasing wall movement may facilitate the OFT tissue to withstand higher ejection pressure and blood volume with development. These findings indicate that OFT remodeling contributes to the effectiveness of blood ejection by increasing wall motion, emphasizing the role of tissue motion in OFT remodeling. Dramatic changes of *in vivo* kinematics of the OFT cushions in the same embryos were also captured along with cardiac motion from HH21 to HH27. The pO and dO cushions start to function independently at HH24 after the intermediate bulging region formation, as demonstrated by cardiac phase lags observed in both cushion closing and opening. It has been shown that changes in endocardial cushion motion were associated with prevalvular tissue elastic rigidity and volume fractions of cells and matrix (67). Our findings may thus indicate a transition in OFT cushion structural composition and properties, which leads to the independent operations of dO and pO instead of passive luminal collapse at early stage HH21.

With *in vivo* longitudinal 4D ultrasound datasets, we then developed quadratic ensemble averaging based algorithms to increase blood-tissue contrast from high-frequency ultrasound, and achieved spatiotemporal synchronization via correlation coefficient-based algorithm. Further, we employed a novel method with custom parameters to automatically track cardiac boundaries. Wall motion boundary data extracted from direct measurement via live scans was applied to surface meshes and registered spatiotemporally with flow conditions as additional constraints, reducing the dependence of fluid boundary conditions while providing more realistic boundary conditions in CFD simulations. This also reduces computational overhead in segmentation and point matching between different models, as displacement functions of surface mesh nodes based on boundary tracking eliminate the need of segmentations at every time frame.

Live-imaging derived moving-domain simulation results showed similar spatial patterns of WSS and velocity, that high WSS results were found in the cushion regions and during OFT lumen shrinking, indicating that the high WSS was due to high blood velocity and smaller lumen diameter and confirmed the earlier finding (28). Statistical results showed that the WSS increased significantly across development in the dO. This is consistent with a previous cohort study using static tissue boundaries (18). However, our peak WSS results are approximately twice as high as those of earlier studies, likely due to our additional cardiac motion boundary conditions. This further emphasizes the interplay between tissue motion and blood flow and the importance of incorporating dynamic boundaries in quantitative hemodynamic studies. High WSS is believed to stimulate vessel remodeling and sculpting of leaflet-shape valves (18, 68). As the aortic and pulmonary valves develop in the dO regions, our data may thus suggest a driving role of increasing WSS in OFT cushion remodeling. Previous studies have demonstrated that low shear stress promotes cell proliferation and cushion growth, whereas high shear stress induces valve shaping and extension in the blood flow direction (15, 17). Our results support these findings using *in vivo* derived *in silico* simulations, that low WSS at earlier developmental stages induces cushion growth, whereas rising WSS across development leads to a switch in morphogenesis at later stages and induces valve shaping. In addition, our OSI results showed peak OSI existed at both sides of the dO and at the middle of inner and outer curvature at HH21 and HH24, same locations as the dO and pO cushions. Reversing flows were recognized as a necessary stimulus to endothelial cells to ensure normal valve morphogenesis in 2-chambered zebrafish embryos without local variations of pO and dO (14). Our results may indicate OSI also plays a role in higher order 4-chambered animal models. Though no statistical significance was found across development, the mean OSI magnitude at HH24 was lower than that at HH21. As previous studies identified oscillatory shear stress as strong stimuli for endocardial proliferation and cushion growth (15, 17), our results may suggest a slowdown in cushion growth from HH21 to HH24. Together with the rising WSS, a transition from cushion growth to cushion compaction and shaping likely happened at around HH24. This is consistent with previous findings that compaction of OFT cushions was observed from HH25 to HH36 (69). Further, higher OSI was observed in the dO than that in the pO at all stages studied, indicating the blood flow to the distal outlets is less aligned in the stream-wise direction than that in the pO. Interestingly, the OSI in dO decreased from HH24 to HH27 in 3 out of 4 embryos, whereas the OSI increased in pO in majority of embryos in the pO (Fig. S3C). The larger oscillation in flow directions in the dO and contrast developmental trends in dO and pO may be associated with the aorticopulmonary septum. The truncus arteriosus septation is believed to proceed in a distal-to-proximal direction toward the ventricles (70). Our simulation results may thus confirm this finding in the hemodynamic perspective that the aorticopulmonary septum developed in the dO first, causing greater flow disturbance with higher OSI and lower WSS in the dO at an earlier stage (HH21). When the dO completed the septation by HH27, OSI in the dO decreased and smaller diameters led to significantly WSS increases, whereas the OSI in the pO began to increase, in consistent with earlier findings that the pO cushions fused by HH30 in the chick embryo (18, 23).

We further quantified the dynamic tissue strains and stresses across development. As expansion strains were calculated, significantly increasing strain values over development were observed as expected, which further confirms our finding that the OFT develops larger expansibility and stretchability over time. An interesting finding was that tissue in pO experiences significantly larger strains than that in the dO at all stages studied. Together with increasing strain values observed in pO across development, these could indicate a transformation in tissue dynamics occurring with pO ventricularization when the proximal OFT myocardium gradually incorporates into the developing right ventricle (71), leading to greater dynamic changes in tissue strains within a cardiac cycle. In addition, lower tissue strains in the dO may be associated with the developing great vessels connecting to the OFT outlet, where limited contraction with heartbeat was observed. Studies on aortic arch morphogenesis modeled the vessel wall as rigid structure since no significant wall motion was observed distal to the OFT (32, 72). This is in line with our observation during ultrasound scan and our tissue mechanics results, that the great vessels distal to the OFT experience little contraction or expansion with heartbeat, leading to lower tissue strain. The dO connecting to those vessels may thus serve as a dampener between the relatively dynamic OFT and relatively fixed great vessels, demonstrating by its significantly smaller strain results to those in the pO. Further, hydrostatic stress results showed that the OFT at HH24 experienced the highest stress rather than at HH27, a result that contrasted with the increase in blood pressure with development. This intriguing result was also accompanied by a large variation in magnitude and in timing among different embryos. These nature variations decreased with the stress magnitudes at HH27, especially in the dO region, suggesting the changes in tissue stress may be related to morphological changes due to the septation of the dO at HH27. In addition, the decrease in hydrostatic stress in dO may also be associated with the growth of arch artery walls. Previous study has reported a marked increase in wall thickness and smooth muscle cells in pharyngeal arch artery (PAA) from HH24 to HH26 (73). Our findings on decreased hydrostatic stress may thus be as results of increased vessel resistance to high blood pressure.

To further delineate the role of hemodynamic forces and tissue mechanics as drivers of OFT remodeling, we correlated WSS and tissue strains with quantified morphological changes in volume and surface area from reconstructed dynamic 3D models. We identified a primary role for WSS in driving the development of stretchability in the OFT, whereas the OFT’s expansive ability was strongly correlated with local tissue strains. In addition, both hemodynamics and tissue strains contribute to the development of tissue stretchability in the dO, while tissue mechanics was also identified as a common driver of both stretchability and expansibility in the dO. As tissue in the dO undergoes complex geometric remodeling such as outlet septation, our results may thus indicate the septation of great vessels requires contributions of both hemodynamic factors and cyclic tissue wall motion. Prior studies of cardiovascular remodeling showed mixed conclusions regarding the relationship between WSS and structural growth. Weak correlations between WSS and vascular size were reported in great vessels in mouse model (74), whereas other studies demonstrated strong correlations between WSS and changes in vascular diameters and cross-sectional areas during PAA remodeling (32, 33, 75). By extending quantitative analysis across both temporal and spatial dimensions, our results refined and advanced the hypothesis that mechanical interactions between blood flow and structural configuration regulate cardiac OFT morphogenesis, and further identified tissue mechanics as an additional and integrated driver during this process. These multi-correlations were not found in the pO, however, suggesting mechanical forces play different roles in the dO and pO development, where other factors contributing to the remodeling of the pO need to be further elucidated. Further, outliers were observed in linear regressions, especially in the dO at HH27, where high WSS magnitudes were not associated with high surface area or volume changes, indicating there may be other factors contributing to tissue mechanics, and hemodynamic force may serve as a driver for other remodeling phenomena.

We then investigated flow helicity to further uncover the hemodynamic driven OFT remodeling. A double-helical flow pattern formed by two counter-rotating flows were observed within the OFT lumen at all developmental stages studied. Spiraling vortex zones due to flow mixing from the left and right side of the OFT were previously reported to start from HH27 in a cohort study using static geometries (18). A similar twisting feature was also reported in a 4D study at HH25, where it was hypothesized to be associated with the aorticopulmonary septation (29). Our results based on high-dimensional simulations extended the observation of this double-helical structure longitudinally across the critical developmental stages when the aorticopulmonary septation occurs, from an earlier stage at 4 days of gestation (HH21) until the septation completed in the dO on D6 (HH27). We noticed the twisting feature was more significant and rapid at earlier stages before dO septation, whereas the two counter-rotating flows interacted more in the dO post-septation at HH27. The flow septation due to counter-rotating flows in the distal OFT at earlier developmental stages might thus be associated with the aorticopulmonary septation that happened between HH24 and HH27, and the helicity in the dO then decreased post-septation. In addition, the axis dividing the two helical flows is in line with the location of aorticopulmonary septum (conotruncal septum) reported by previous anatomy studies (76–78). Our findings may thus relate aorticopulmonary septation to the observed blood flow pattern, supporting the hypothesis by Ho et al. that septum develops in between the two helical flows as a result of flow septation due to counter-rotating flows with a longitudinal study tracking morphing in the same embryos. Intriguingly, the magnitudes of the normalized helicity iso-surfaces close to the tissue are 0.05 and -0.05, which physically means that the angle between the forward velocity vector and the vorticity vector is around 87.13 degrees. This indicates that flow direction shifts as small as ∼3 degrees may still be sensible to the endocardial cells and trigger mechanotransduction, suggesting an intriguingly lower threshold in cellular mechanosensing than physiological conditions and previously identified (79, 80). In addition, this double-helical feature may contribute to the balance of systemic and pulmonary circulations at earlier developmental stages prior to aorticopulmonary septation. Such features were also observed in lower species such as frog, where a valve-like spiral structure was observed in the conus arteriosus and is believed to be responsible to direct oxidized and non-oxidized blood flow in a single-outlet ventricle (81, 82). Our findings may thus indicate an evolutionary legacy that can be further correlated to CHDs with single ventricle (83). Together, our findings further emphasized the driving role of hemodynamic forces in cardiac development and OFT remodeling.

In conclusion, this study provides longitudinal live image-derived moving-domain CFD simulations of a developing OFT in a four-chambered animal model within a full cardiac cycle across critical valvulogenesis developmental stages in the same embryos. By co-analyzing the quantified hemodynamic indices and local tissue stresses with the morphing OFT, we identified hemodynamic force and wall motion as drivers of local tissue development and important stimuli for OFT remodeling. We employed 4D live imaging with CFD simulations to achieve spatiotemporal mappings of hemodynamics and tissue strains. Wall motion derived from *in vivo* ultrasound scans was implemented as additional boundary conditions to increase accuracy and realness. This extended our understanding in hemodynamics when co-analyzed to earlier studies using static boundaries. In addition, this novel approach provides opportunities to rapidly analyze dynamic tissue mechanics by directly using surface nodal displacement data extracted from boundary tracking without additional fluid-structure interaction solver needed. While the current application focuses on OFT remodeling, this approach can be applied to any embryonic cardiac structure. A combination of this technique with experimental perturbation or molecular manipulation will significantly enhance our understanding of cardiac morphogenesis and CHD pathogenesis.

## Acknowledgements

We thank the Cornell BRC Imaging facility, Fred Jia (simulation), Shuofei Sun (ultrasound), and members of Butcher lab for helpful discussion.

## Author contributions

Conceptualization: G.D., J.T.B.; Methodology: G.D., J.T.B., H.L., M.E.; Software: G.D., J.R., S.K., M.E.D., H.L., M.E.; Validation: G.D., J.R., S.K., M.E.D.; Formal analysis: G.D.; Investigation: G.D., J.R.; Resources: J.T.B., M.E.; Data curation: G.D., J.R.; Writing - original draft: G.D.; Writing - review & editing: J.T.B.; Visualization: G.D.; Supervision: J.T.B, M.E.; Project administration: J.T.B., M.E.; Funding acquisition: J.T.B.

## References

1. K. Courchaine, M. J. Gray, K. Beel, K. Thornburg, S. Rugonyi, 4-D Computational Modeling of Cardiac Outflow Tract Hemodynamics over Looping Developmental Stages in Chicken Embryos. J Cardiovasc Dev Dis 6, 11 (2019).

2. R. H. Anderson, S. Mori, D. E. Spicer, N. A. Brown, T. J. Mohun, Development and Morphology of the Ventricular Outflow Tracts. World J Pediatr Congenit Heart Surg 7, 561–577 (2016).

3. R. E. Poelmann, et al., Ventricular Septation and Outflow Tract Development in Crocodilians Result in Two Aortas with Bicuspid Semilunar Valves. J Cardiovasc Dev Dis 8, 132 (2021).

4. Z. Neeb, J. D. Lajiness, E. Bolanis, S. J. Conway, Cardiac outflow tract anomalies. WIREs Developmental Biology 2, 499–530 (2013).

5. S. Stefanovic, H. C. Etchevers, S. Zaffran, Outflow Tract Formation—Embryonic Origins of Conotruncal Congenital Heart Disease. J Cardiovasc Dev Dis 8, 42 (2021).

6. M. E. Pierpont, et al., Genetic Basis for Congenital Heart Disease: Revisited: A Scientific Statement From the American Heart Association. Circulation 138 (2018).

7. W. Wu, J. He, X. Shao, Incidence and mortality trend of congenital heart disease at the global, regional, and national level, 1990–2017. Medicine 99, e20593 (2020).

8. J. I. E. Hoffman, S. Kaplan, The incidence of congenital heart disease. J Am Coll Cardiol 39, 1890–1900 (2002).

9. B. M. Jepson, et al., Pregnancy loss in major fetal congenital heart disease: incidence, risk factors and timing. Ultrasound in Obstetrics & Gynecology 62, 75–87 (2023).

10. S. N. Nees, W. K. Chung, Genetic Basis of Human Congenital Heart Disease. Cold Spring Harb Perspect Biol 12, a036749 (2020).

11. H. E. Salman, et al., Effect of left atrial ligation-driven altered inflow hemodynamics on embryonic heart development: clues for prenatal progression of hypoplastic left heart syndrome. Biomech Model Mechanobiol 20, 733–750 (2021).

12. H. C. Yalcin, et al., Two-photon microscopy-guided femtosecond-laser photoablation of avian cardiogenesis: noninvasive creation of localized heart defects. American Journal of Physiology-Heart and Circulatory Physiology 299, H1728–H1735 (2010).

13. H. Fukui, et al., Bioelectric signaling and the control of cardiac cell identity in response to mechanical forces. Science (1979) 374, 351–354 (2021).

14. J. Vermot, et al., Reversing Blood Flows Act through klf2a to Ensure Normal Valvulogenesis in the Developing Heart. PLoS Biol 7, e1000246 (2009).

15. M. Wang, et al., Shear and hydrostatic stress regulate fetal heart valve remodeling through YAP-mediated mechanotransduction. Elife 12 (2023).

16. L. M. Goddard, et al., Hemodynamic Forces Sculpt Developing Heart Valves through a KLF2-WNT9B Paracrine Signaling Axis. Dev Cell 43, 274-289.e5 (2017).

17. D. H. Pham, C. R. Dai, B. Y. Lin, J. T. Butcher, Local fluid shear stress operates a molecular switch to drive fetal semilunar valve extension. Developmental Dynamics 251, 481–497 (2022).

18. K. N. Bharadwaj, C. Spitz, A. Shekhar, H. C. Yalcin, J. T. Butcher, Computational Fluid Dynamics of Developing Avian Outflow Tract Heart Valves. Ann Biomed Eng 40, 2212– 2227 (2012).

19. V. Menon, J. Eberth, R. Goodwin, J. Potts, Altered Hemodynamics in the Embryonic Heart Affects Outflow Valve Development. J Cardiovasc Dev Dis 2, 108–124 (2015).

20. S. Goenezen, V. K. Chivukula, M. Midgett, L. Phan, S. Rugonyi, 4D subject-specific inverse modeling of the chick embryonic heart outflow tract hemodynamics. Biomech Model Mechanobiol 15, 723–743 (2016).

21. A. Liu, et al., Biomechanics of the Chick Embryonic Heart Outflow Tract at HH18 Using 4D Optical Coherence Tomography Imaging and Computational Modeling. PLoS One 7, e40869 (2012).

22. V. Chivukula, S. Goenezen, A. Liu, S. Rugonyi, Effect of Outflow Tract Banding on Embryonic Cardiac Hemodynamics. J Cardiovasc Dev Dis 3, 1 (2015).

23. J. G. Wittig, A. Münsterberg, The Chicken as a Model Organism to Study Heart Development. Cold Spring Harb Perspect Biol 12, a037218 (2020).

24. V. Messerschmidt, et al., Light-sheet Fluorescence Microscopy to Capture 4-Dimensional Images of the Effects of Modulating Shear Stress on the Developing Zebrafish Heart. Journal of Visualized Experiments (2018). 10.3791/57763.

25. J. Lee, et al., Moving Domain Computational Fluid Dynamics to Interface with an Embryonic Model of Cardiac Morphogenesis. PLoS One 8, e72924 (2013).

26. A. S. Forouhar, et al., The Embryonic Vertebrate Heart Tube Is a Dynamic Suction Pump. Science (1979) 312, 751–753 (2006).

27. V. Vedula, et al., A method to quantify mechanobiologic forces during zebrafish cardiac development using 4-D light sheet imaging and computational modeling. PLoS Comput Biol 13, e1005828 (2017).

28. S. Ho, W. X. Chan, N. Phan-Thien, C. H. Yap, Organ Dynamics and Hemodynamic of the Whole HH25 Avian Embryonic Heart, Revealed by Ultrasound Biomicroscopy, Boundary Tracking, and Flow Simulations. Sci Rep 9, 18072 (2019).

29. S. Ho, W. X. Chan, S. Rajesh, N. Phan-Thien, C. H. Yap, Fluid dynamics and forces in the HH25 avian embryonic outflow tract. Biomech Model Mechanobiol 18, 1123–1137 (2019).

30. S. Ho, G. X. Y. Tan, T. J. Foo, N. Phan-Thien, C. H. Yap, Organ Dynamics and Fluid Dynamics of the HH25 Chick Embryonic Cardiac Ventricle as Revealed by a Novel 4D High-Frequency Ultrasound Imaging Technique and Computational Flow Simulations. Ann Biomed Eng 45, 2309–2323 (2017).

31. D. Zhang, S. E. Lindsey, Rapid Whole-Mount High-Resolution Imaging of Small Animal Vasculature for Quantitative Studies. Journal of Visualized Experiments (2025). 10.3791/68206.

32. S. E. Lindsey, J. T. Butcher, I. E. Vignon-Clementel, Cohort-based multiscale analysis of hemodynamic-driven growth and remodeling of the embryonic pharyngeal arch arteries. Development 145 (2018).

33. S. E. Lindsey, I. E. Vignon-Clementel, J. T. Butcher, Assessing Early Cardiac Outflow Tract Adaptive Responses Through Combined Experimental-Computational Manipulations. Ann Biomed Eng 49, 3227–3242 (2021).

34. M. Midgett, V. K. Chivukula, C. Dorn, S. Wallace, S. Rugonyi, Blood flow through the embryonic heart outflow tract during cardiac looping in HH13–HH18 chicken embryos. J R Soc Interface 12, 20150652 (2015).

35. K. Balachandran, et al., Cyclic strain induces dual-mode endothelial-mesenchymal transformation of the cardiac valve. Proceedings of the National Academy of Sciences 108, 19943–19948 (2011).

36. B. J. Damon, et al., Patterns of muscular strain in the embryonic heart wall. Developmental Dynamics 238, 1535–1546 (2009).

37. S. E. Lindsey, Mechanical regulation of cardiac development. Front Physiol 5 (2014).

38. S. Rugonyi, Strain-Induced Tissue Growth Laws: Applications to Embryonic Cardiovascular Development. J Appl Mech Eng s11 (2013).

39. H. C. Yalcin, A. Shekhar, A. A. Rane, J. T. Butcher, An ex-ovo Chicken Embryo Culture System Suitable for Imaging and Microsurgery Applications. Journal of Visualized Experiments 2154 (2010). 10.3791/2154.

40. A. Dittrich, M. M. Thygesen, H. Lauridsen, 2D and 3D Echocardiography in the Axolotl (Ambystoma Mexicanum). Journal of Visualized Experiments (2018). 10.3791/57089.

41. A. Fedorov, et al., 3D Slicer as an image computing platform for the Quantitative Imaging Network. Magn Reson Imaging 30, 1323–1341 (2012).

42. C. Pinter, A. Lasso, G. Fichtinger, Polymorph segmentation representation for medical image computing. Comput Methods Programs Biomed 171, 19–26 (2019).

43. M. Esmaily-Moghadam, Y. Bazilevs, A. L. Marsden, A bi-partitioned iterative algorithm for solving linear systems arising from incompressible flow problems. Comput Methods Appl Mech Eng 286, 40–62 (2015).

44. M. Esmaily-Moghadam, Y. Bazilevs, A. L. Marsden, A new preconditioning technique for implicitly coupled multidomain simulations with applications to hemodynamics. Comput Mech 52, 1141–1152 (2013).

45. M. Esmaily-Moghadam, I. E. Vignon-Clementel, R. Figliola, A. L. Marsden, A modular numerical method for implicit 0D/3D coupling in cardiovascular finite element simulations. J Comput Phys 244, 63–79 (2013).

46. P. Cignoni, et al., MeshLab: an open-source mesh processing tool. Eurographics Italian chapter conference 2008, 129–136 (2008).

47. H. Si, TetGen, a Delaunay-Based Quality Tetrahedral Mesh Generator. ACM Transactions on Mathematical Software 41, 1–36 (2015).

48. A. I. Hassaballah, M. A. Hassan, A. N. Mardi, M. Hamdi, An Inverse Finite Element Method for Determining the Tissue Compressibility of Human Left Ventricular Wall during the Cardiac Cycle. PLoS One 8, e82703 (2013).

49. E. B. Clark, et al., Ventricular function and morphology in chick embryo from stages 18 to 29. American Journal of Physiology-Heart and Circulatory Physiology 250, H407– H413 (1986).

50. E. B. Clark, et al., Effect of increased pressure on ventricular growth in stage 21 chick embryos. American Journal of Physiology-Heart and Circulatory Physiology 257, H55– H61 (1989).

51. N. Hu, D. M. Connuck, B. B. Keller, E. B. Clark, Diastolic Filling Characteristics in the Stage 12 to 27 Chick Embryo Ventricle. Pediatr Res 29, 334–337 (1991).

52. N. Hu, E. B. Clark, Hemodynamics of the stage 12 to stage 29 chick embryo. Circ Res 65, 1665–1670 (1989).

53. J. P. Ahrens, B. Geveci, C. C. Law, ParaView: An End-User Tool for Large-Data Visualization in The Visualization Handbook, (2005).

54. H. Wickham, ggplot2: Elegant Graphics for Data Analysis (Springer-Verlag New York, 2016).

55. B. Cuneo, S. Hughes, D. W. Benson, Heart rate perturbation in the stage 17-27 chick embryo: effect on stroke volume and aortic flow. American Journal of Physiology-Heart and Circulatory Physiology 264, H755–H759 (1993).

56. R. H. Anderson, et al., Development of the arterial roots and ventricular outflow tracts. J Anat (2023). 10.1111/joa.13973.

57. R. H. Anderson, S. Mori, D. E. Spicer, N. A. Brown, T. J. Mohun, Development and Morphology of the Ventricular Outflow Tracts. World J Pediatr Congenit Heart Surg 7, 561–577 (2016).

58. R. Lorenz, et al., 4D flow magnetic resonance imaging in bicuspid aortic valve disease demonstrates altered distribution of aortic blood flow helicity. Magn Reson Med 71, 1542–1553 (2014).

59. H. C. Yalcin, A. Shekhar, T. C. McQuinn, J. T. Butcher, Hemodynamic patterning of the avian atrioventricular valve. Developmental Dynamics 240, 23–35 (2011).

60. M. M. Ossa Galvis, R. T. Bhakta, A. Tarmahomed, M. D. Mendez, Cyanotic Heart Disease (2025).

61. M. C. Snoep, et al., Factors related to fetal demise in cases with congenital heart defects. Am J Obstet Gynecol MFM 5, 101023 (2023).

62. L. Song, et al., Myocardial Smad4 Is Essential for Cardiogenesis in Mouse Embryos. Circ Res 101, 277–285 (2007).

63. S. J. Conway, A. Kruzynska‐Frejtag, P. L. Kneer, M. Machnicki, S. V. Koushik, What cardiovascular defect does my prenatal mouse mutant have, and why? genesis 35, 1– 21 (2003).

64. M. C. Mak, et al., Embryonic Lethality in Mice Lacking the Nuclear Factor of Activated T Cells 5 Protein Due to Impaired Cardiac Development and Function. PLoS One 6, e19186 (2011).

65. L. M. Goddard, et al., Hemodynamic Forces Sculpt Developing Heart Valves through a KLF2-WNT9B Paracrine Signaling Axis. Dev Cell 43, 274-289.e5 (2017).

66. B. J. Martinsen, Reference guide to the stages of chick heart embryology. Developmental Dynamics 233, 1217–1237 (2005).

67. J. T. Butcher, T. C. McQuinn, D. Sedmera, D. Turner, R. R. Markwald, Transitions in Early Embryonic Atrioventricular Valvular Function Correspond With Changes in Cushion Biomechanics That Are Predictable by Tissue Composition. Circ Res 100, 1503–1511 (2007).

68. S. Chen, H. Zhang, Q. Hou, Y. Zhang, A. Qiao, Multiscale Modeling of Vascular Remodeling Induced by Wall Shear Stress. Front Physiol 12 (2022).

69. D. Bassen, et al., Hydrostatic mechanical stress regulates growth and maturation of the atrioventricular valve. Development 148 (2021).

70. T. J. Mohun, R. H. Anderson, 3D Anatomy of the Developing Heart: Understanding Ventricular Septation. Cold Spring Harb Perspect Biol 12, a037465 (2020).

71. M. S. Rana, et al., Trabeculated Right Ventricular Free Wall in the Chicken Heart Forms by Ventricularization of the Myocardium Initially Forming the Outflow Tract. Circ Res 100, 1000–1007 (2007).

72. Y. Wang, et al., Aortic Arch Morphogenesis and Flow Modeling in the Chick Embryo. Ann Biomed Eng 37, 1069–1081 (2009).

73. J. Ryvlin, S. E. Lindsey, J. T. Butcher, Systematic Analysis of the Smooth Muscle Wall Phenotype of the Pharyngeal Arch Arteries During Their Reorganization into the Great Vessels and Its Association with Hemodynamics. Anat Rec 302, 153–162 (2019).

74. C. H. Yap, X. Liu, K. Pekkan, Characterizaton of the Vessel Geometry, Flow Mechanics and Wall Shear Stress in the Great Arteries of Wildtype Prenatal Mouse. PLoS One 9, e86878 (2014).

75. W. J. Kowalski, et al., Critical Transitions in Early Embryonic Aortic Arch Patterning and Hemodynamics. PLoS One 8, e60271 (2013).

76. D. A. Goor, R. Dische, C. W. Lillehei, The Conotruncus. Circulation 46, 375–384 (1972).

77. G. del Monte-Nieto, R. P. Harvey, “Embryology of the Heart” in Skin and the Heart, (Springer International Publishing, 2021), pp. 11–30.

78. R. H. Anderson, Development of the heart: (3) Formation of the ventricular outflow tracts, arterial valves, and intrapericardial arterial trunks. Heart 89, 1110–1118 (2003).

79. C. A. Dessalles, C. Leclech, A. Castagnino, A. I. Barakat, Integration of substrate- and flow-derived stresses in endothelial cell mechanobiology. Commun Biol 4, 764 (2021).

80. I. Andreu, et al., The force loading rate drives cell mechanosensing through both reinforcement and cytoskeletal softening. Nat Commun 12, 4229 (2021).

81. F. J. Haberich, THE FUNCTIONAL SEPARATION OF VENOUS AND ARTERIAL BLOOD IN THE UNIVENTRICULAR FROG HEART. Ann N Y Acad Sci 127, 459–476 (1965).

82. A. F. Corno, et al., The Secrets of the Frogs Heart. Pediatr Cardiol 43, 1471–1480 (2022).

83. S. L. Meyer, et al., Intracardiac anatomical relationships and potential for streaming in double inlet left ventricles. PLoS One 12, e0188048 (2017).

