## Supplemental Figure 1-7, Supplemental Video 1-3 for "Longitudinal Live Imaging Derived 4D Hemodynamics and Dynamic Tissue Mechanics Across Outflow Tract Morphogenesis": supplemental files_longitudinal_4D_BiorXiv.pdf

### Supplemental Figures

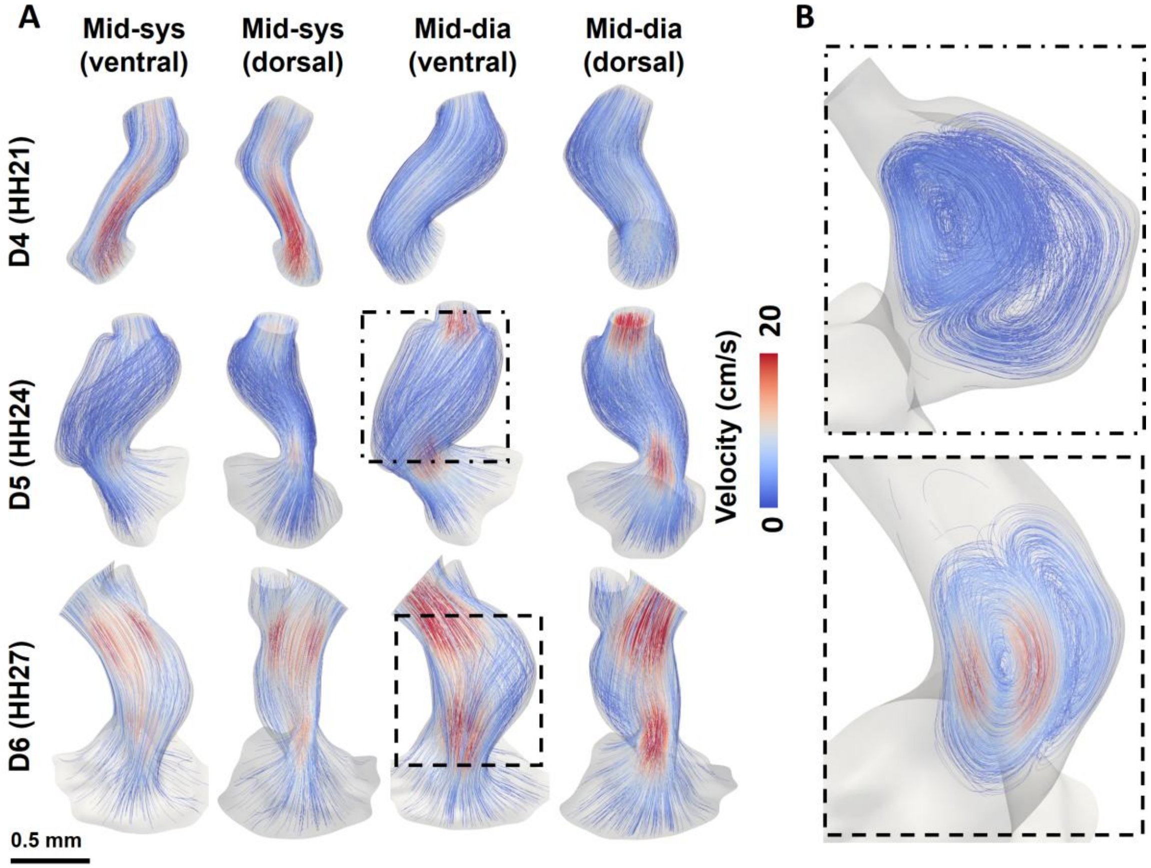


**Figure S1: Time-varying streamline within OFT lumen. A)** Time- and space-varying streamlines contoured with velocity magnitudes at different phases of cardiac cycle over development. Figures show the 3D renderings of the velocity streamlines within the dynamic models of Embryo 1 at different developmental stages at mid-systole and mid-diastole. Higher velocity magnitudes were observed at the proximal cushions and distal cushions during systole after cushion opening. **B)** Vortex flow in the intermediate OFT region during cushion shut.

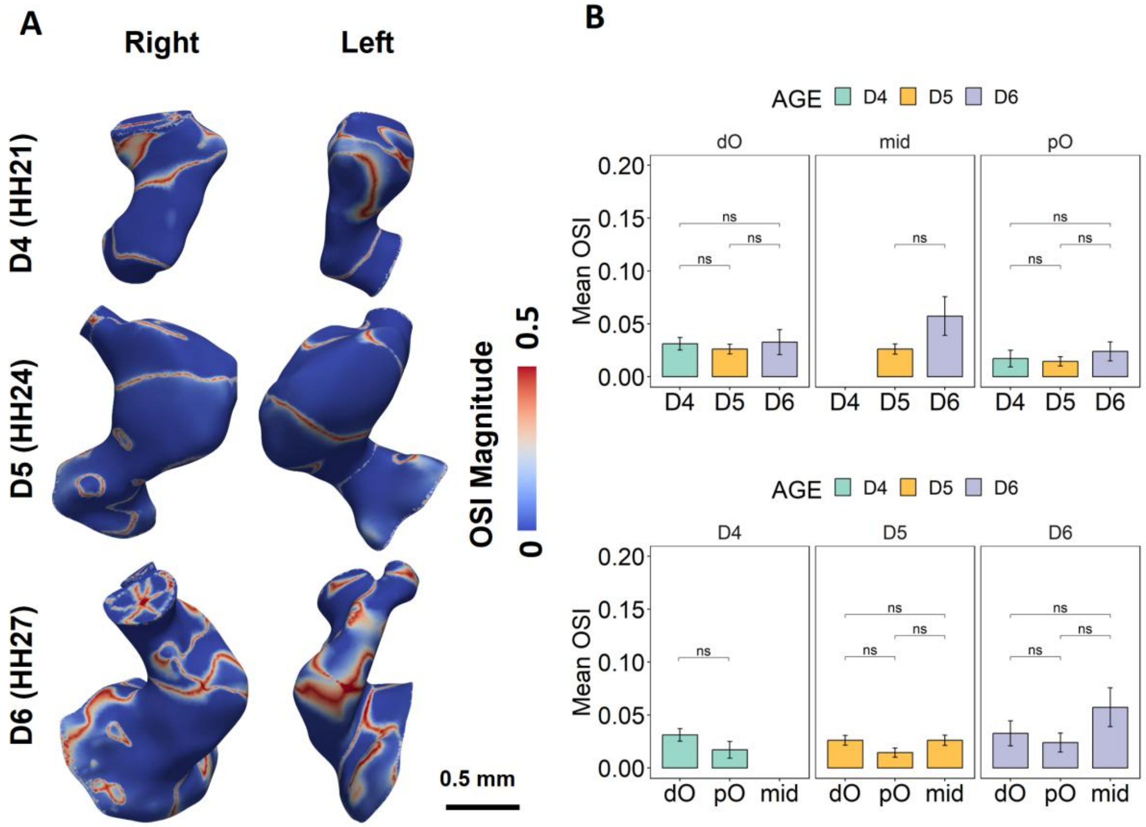


**Figure S2: Oscillatory shear index in the OFT lumen across development. A)** Models of Embryo 1 at the segmentation time frame were used to show the spatial variation of OSI across development. Higher OSI in the intermediate OFT region at HH24 and HH27. **B)** No difference in regional mean OSI magnitudes over time or among regions.

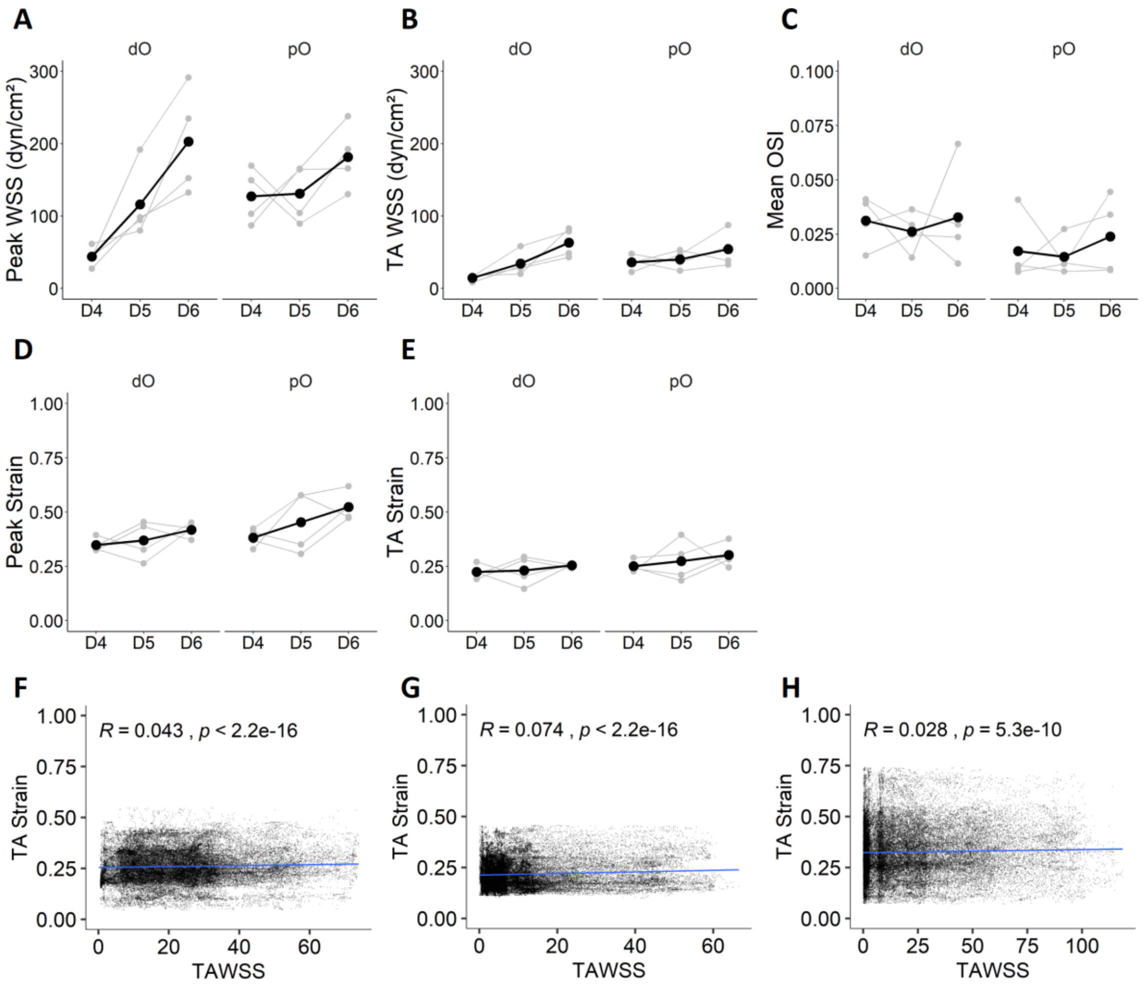


**Figure S3: Developmental trends of A) peak WSS, B) time-averaged WSS, C) OSI, D) peak strain, and E) time-averaged strain in 4 embryos.** Data points from the same embryo were linked with black lines across development. **No spatial correlations between strain features and WSS at F) HH21, G) HH24, and H) HH27.** Data points represent strain and TAWSS magnitudes of the same elements.

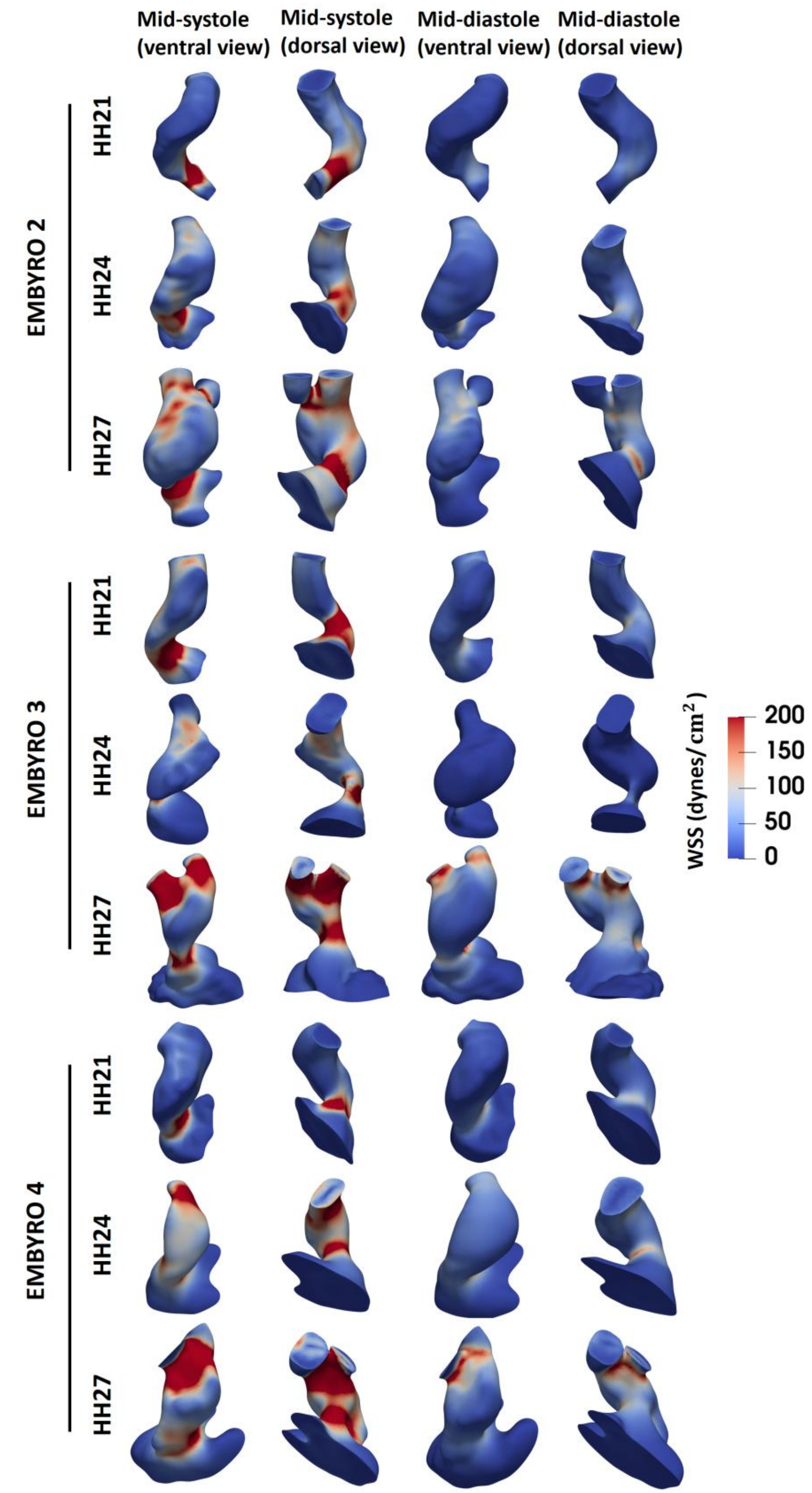


**Figure S4:** Time- and space-varying WSS magnitudes at different phases of cardiac cycle over development for Embryo 2,3, and 4.

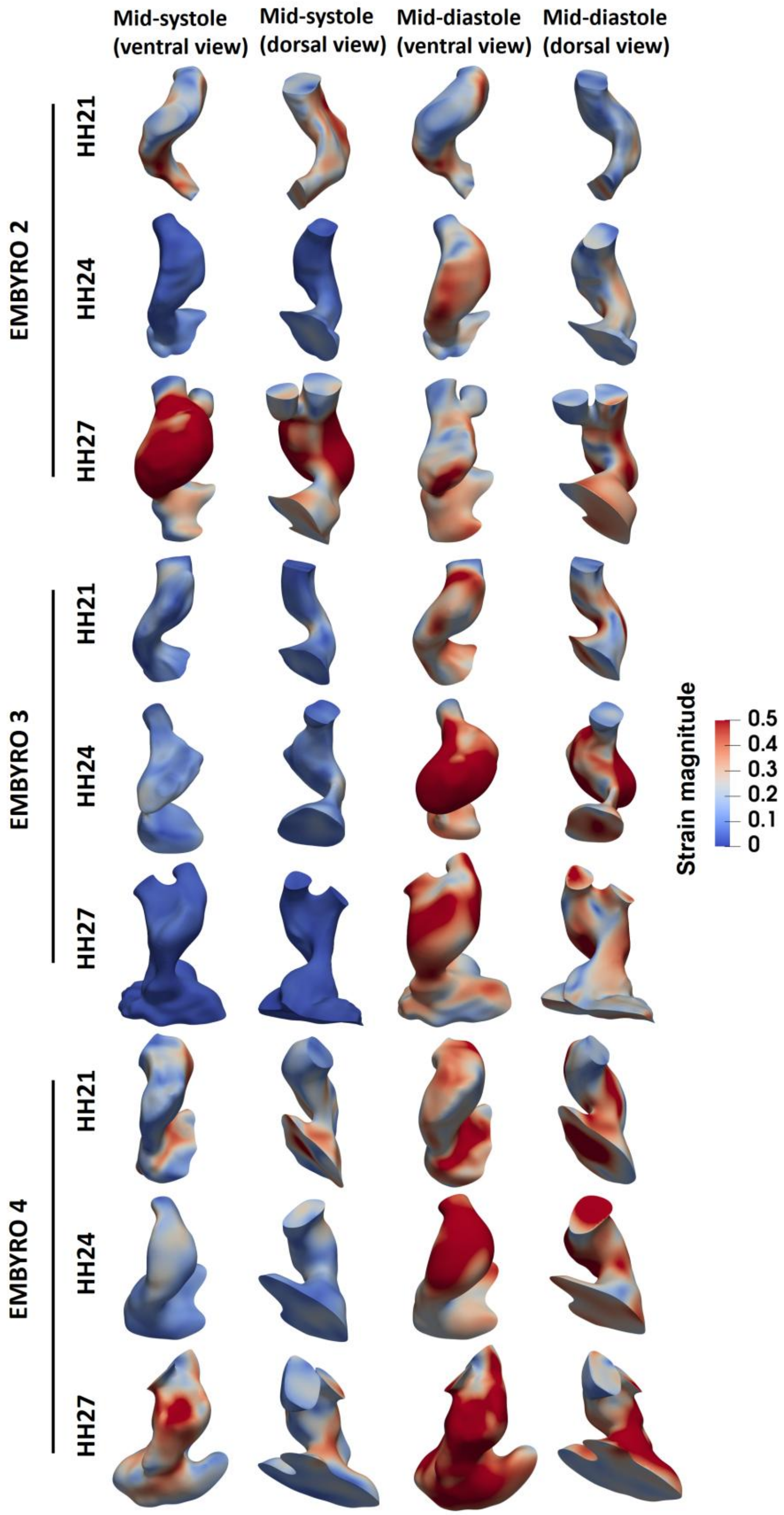


**Figure S5:** Time- and space-varying strain magnitudes at different phases of cardiac cycle over development for Embryo 2,3, and 4.

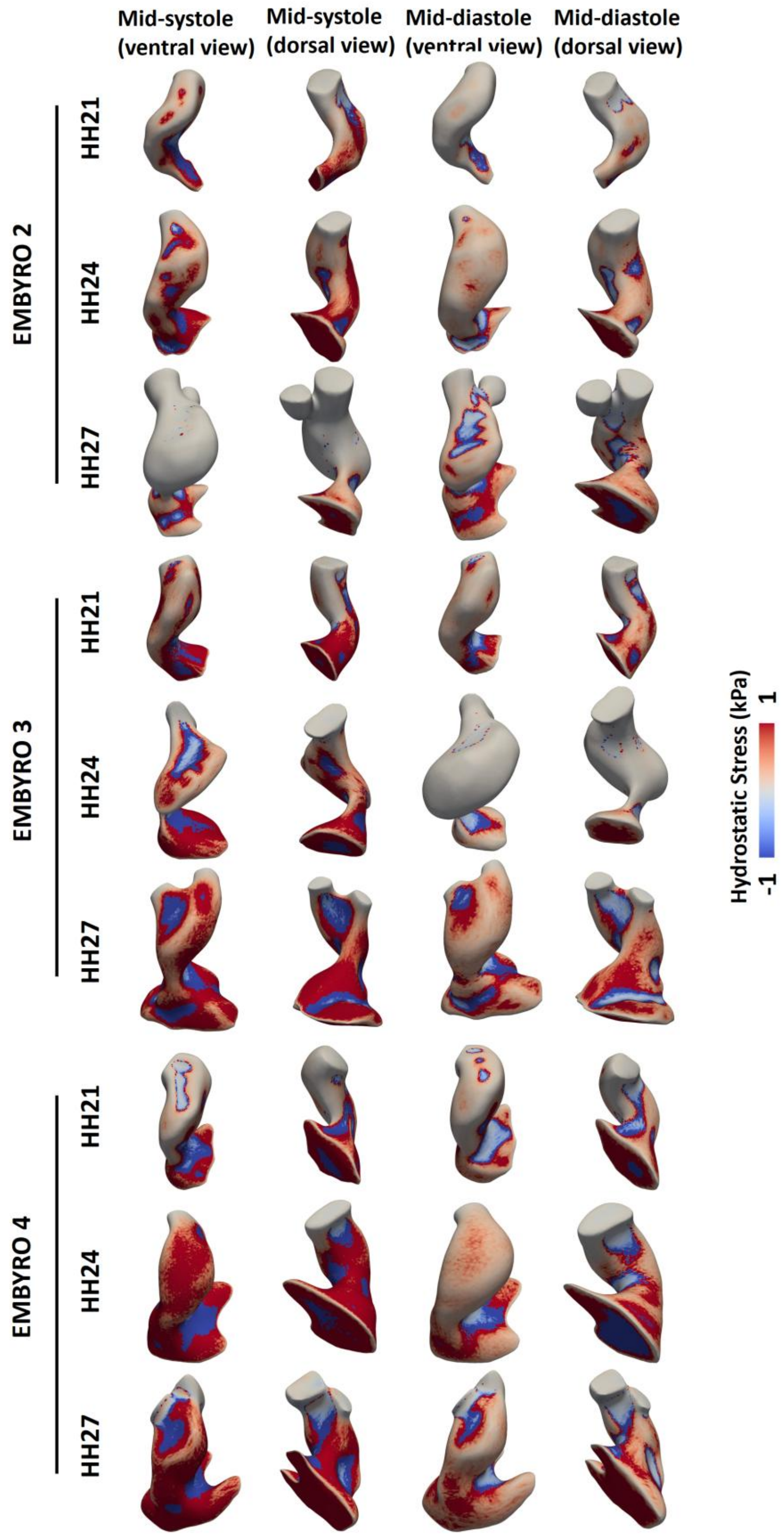


**Figure S6:** Time- and space-varying hydrostatic wall stress at different phases of cardiac cycle over development for Embryo 2,3, and 4.

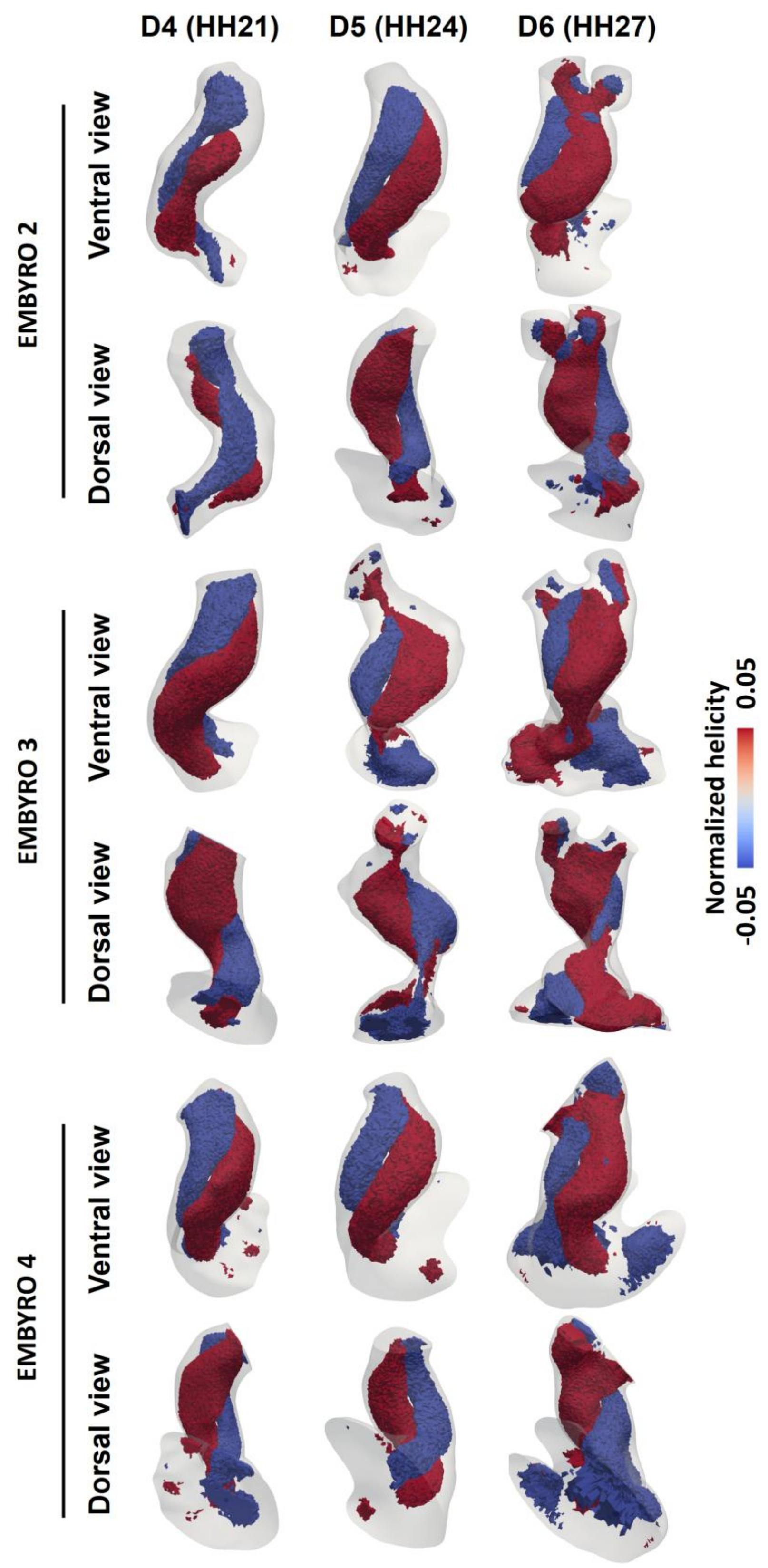


**Figure S7:** Double-helical flow pattern observed during blood ejection after dO cushions opening, shown in the models of Embryo 2, 3, 4 at different developmental stages.

### Supplemental Video Legends

**Video S1:** Ultrasound cine images showed transition in cushion motions across development at **A)** HH21, **B)** HH24, **C)** HH27. * indicates pO and dO locations.

**Video S2:** Time- and space-varying WSS magnitudes within a full cardiac cycle over development for Embryo 1 at **A)** HH21, **B)** HH24, **C)** HH27. Arrows indicate blood flow directions. V, ventricle.

**Video S3:** Time- and space-varying expansive strain from minimum volume within a full cardiac cycle over development for Embryo 1 at **A)** HH21, **B)** HH24, **C)** HH27. Arrows indicate blood flow directions. V, ventricle.
